# *Legionella pneumophila* macrophage infectivity potentiator protein appendage domains modulate protein dynamics and inhibitor binding

**DOI:** 10.1101/2023.04.24.538046

**Authors:** C. Wiedemann, J.J. Whittaker, V.H. Pérez Carrillo, B. Goretzki, M. Dajka, F. Tebbe, J.-M. Harder, P. Krajczy, B. Joseph, F. Hausch, A. Guskov, U.A. Hellmich

## Abstract

Macrophage infectivity potentiator (MIP) proteins are widespread in human pathogens including *Legionella pneumophila*, the causative agent of Legionnaires’ disease and protozoans such as *Trypanosoma cruzi*. All MIP proteins contain a FKBP (FK506 binding protein)-like prolyl-*cis/trans*- isomerase domain that hence presents an attractive drug target. Some MIPs such as the *Legionella pneumophila* protein (*Lp*MIP) have additional appendage domains of mostly unknown function. In full-length, homodimeric *Lp*MIP, the N-terminal dimerization domain is linked to the FKBP-like domain via a long, free-standing stalk helix. Combining X-ray crystallography, NMR and EPR spectroscopy and SAXS, we elucidated the importance of the stalk helix for protein dynamics and inhibitor binding to the FKBP-like domain and bidirectional crosstalk between the different protein regions. The first comparison of a microbial MIP and a human FKBP in complex with the same synthetic inhibitor was made possible by high-resolution structures of *Lp*MIP with a [4.3.1]-aza-bicyclic sulfonamide and provides a basis for designing pathogen-selective inhibitors. Through stereospecific methylation, the affinity of inhibitors to *L. pneumophila* and *T. cruzi* MIP was greatly improved. The resulting X-ray inhibitor-complex structures of *Lp*MIP and *Tc*MIP at 1.49 and 1.34 Å, respectively, provide a starting point for developing potent inhibitors against MIPs from multiple pathogenic microorganisms.

## Introduction

Bacterial parasitism is a wide-spread phenomenon and a serious health concern [1]. Approximately half of all identified *Legionella* species are associated with human disease, but most human legionellosis are caused by *Legionella pneumophila* [2]. In their natural fresh water reservoir habitat, these facultative intracellular gram-negative bacteria infect protozoa, where, protected from harsh environmental conditions, they find optimal conditions for intracellular replication while benefiting from the nutrient supply provided by the host [3]. After aspiration of contaminated water from e.g. air conditioners or hot water cisterns, *L. pneumophila* can also invade alveolar macrophages in the human lung thereby mimicking the infection of its native amoebal host [2,4,5]. This may result in severe infections such as Legionnaires’ disease or the more benign Pontiac disease [2,4]. Although *Legionella* infections can be treated with antibiotics, Legionnaires’ disease nonetheless has a mortality rate of ∼10%, which is likely even higher in older or immunocompromised patients [6].

To promote uptake into a host cell, *L. pneumophila* relies on a number of proteins, including MIP (Macrophage infectivity potentiator), the first identified *L. pneumophila* virulence factor [7–9]. *Legionella pneumophila* MIP (*Lp*MIP) improves the environmental fitness of the bacterium and facilitates the progression of the early stages of the intracellular infection cycle [9–11]. Genetic deletion of *Lp*MIP results in a reduced intracellular replication rate [9,12].

*Lp*MIP is a homodimeric protein consisting of an N-terminal dimerization domain, a 65 Å long, free- standing α-helix, the “stalk helix”, and a C-terminal peptidyl prolyl-*cis/trans*-isomerase (PPIase) domain [13–15]. Structurally, the PPIase domain belongs to the FK506-binding proteins (FKBPs) named after their interaction with the natural product macrolide lactone FK506 [16,17]. In FKBPs, an amphipathic five-stranded β-sheet wraps around an α-helix thus forming a hydrophobic cavity that binds substrates and inhibitors [18]. Although the molecular mechanism of *Lp*MIP action in infection and its molecular target(s) remain unclear, it was implicated in host collagen interaction and subsequent epithelial barrier transmigration [19,20]. Nonetheless, the interaction between *Lp*MIP and collagen could not be mapped in detail, and instead of using classic chemical shift perturbations (CSP), NMR (nuclear magnetic resonance) spectroscopic PREs (paramagnetic relaxation enhancement) of spin-labeled collagen peptides had to be used to detect binding to *Lp*MIP [19], suggesting weak binding affinities. In contrast, unambiguous binding site mapping to *Lp*MIP has been shown by NMR CSP for rapamycin, a macrolide which also inhibits human FKBPs [21].

MIP proteins are widely expressed in many other human pathogenic microorganisms such as *Chlamydia spp.* [22], *Neisseria gonorrhoeae* [23], the entero-pathogen *Salmonella typhimurium* [24]*, Pseudomonas aeruginosa* [25], and intracellular parasitic protozoans such as *Trypanosoma cruzi*, the causative agent of Chagas disease in South and Central America [26–28]. Hence, the PPIase domains of MIP proteins are attractive antimicrobial and antiparasitic drug targets [29], however their shallow ligand binding pocket and similarity to human FKBPs render selective drug design challenging [30,31]. No structures of a *Legionella* MIP with a synthetic inhibitor are available to date and, in the absence of a high-resolution structure of a microbial MIP and human FKBP MIP in complex with the same synthetic inhibitor, no side-by-side structural comparison is currently possible.

Limited structural information of *Lp*MIP is available, with only a crystal structure of the *apo* full-length homodimer (PDB: 1FD9) [14] and the NMR solution structures of an *apo* and rapamycin-bound truncation mutant (PDB: 2UZ5, 2VCD) [21]. This construct, *Lp*MIP^77-213^, comprises the C-terminal half of the stalk helix followed by the FKBP-like domain and thus resembles the architecture of the constitutively monomeric *T. cruzi* MIP protein [26]. Other pathogens such as *Burkholderia pseudomallei,* the bacterium causing melioidosis, express even more minimalistic MIP proteins, lacking both dimerization domain and the complete stalk helix [32,33].

The role of MIP appendage domains, or the consequences of their (partial) absence, remains unclear. However, homodimeric, full-length MIP from *Legionella pneuomophila* presents a unique opportunity to explore the role of these domains in conformational flexibility and inhibitor binding. Here, we combined X-ray crystallography, small angle X-ray scattering (SAXS), nuclear magnetic resonance (NMR) and electron paramagnetic resonance (EPR) spectroscopy to uncover the importance of the *Lp*MIP stalk helix for the protein’s functional dynamics and to identify similarities and differences in inhibitor binding among MIP proteins from various human pathogenic microorganisms and human FKBPs.

## Results

### Structural dynamics of full-length LpMIP and consequences of inhibitor binding

Comparing our crystal structure of homodimeric full-length *Lp*MIP with improved resolution (1.71 Å, PDB: 8BJC) to the previously published one (2.41 Å, PDB: 1FD9 [14]), revealed a ∼18° splay between the stalk helices in the two structures (Fig. 1A, B). The higher resolution of our electron density map allowed unambiguous placement and assignment of all stalk helix residues (Fig. 1C, Table S1). Furthermore, the stalk helix is not involved in crystal contacts suggesting that intrinsic conformational heterogeneity is responsible for the observed differences between the two structures.

**Fig. 1:**
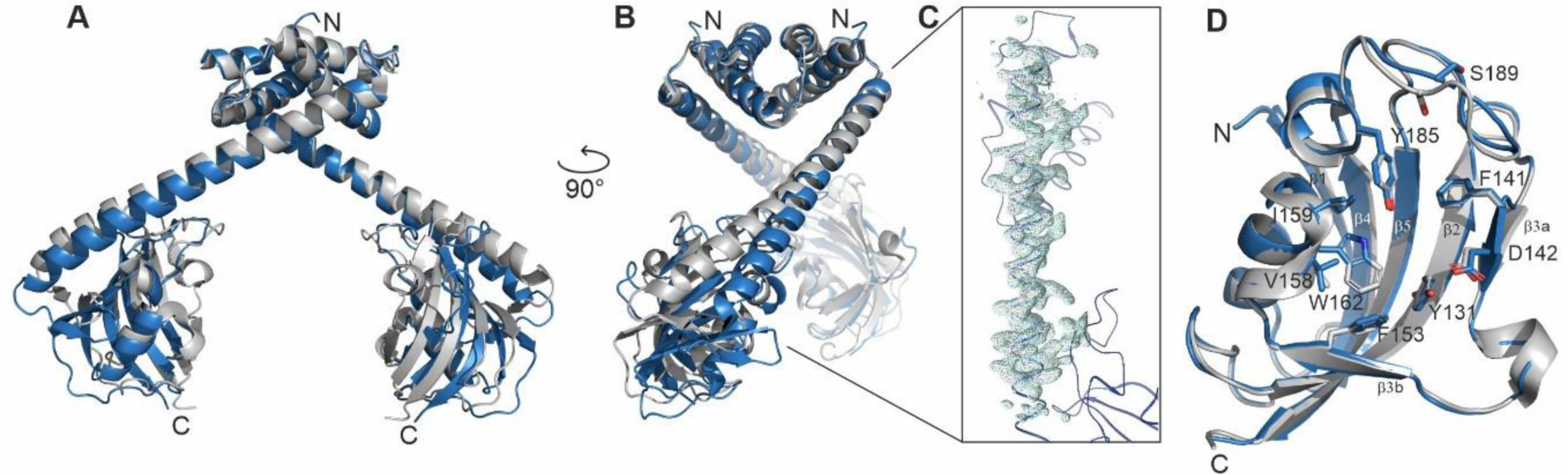
Comparison of full-length *Lp*MIP structures reveal stalk helix splaying. (**A, B**) Overlay of the N-terminal dimerization domains of the two currently available *Lp*MIP^1-213^ structures (PDB: 1FD9 at 2.41 Å, grey; PDB: 8BJC at 1.71 Å, blue) shows ∼18° stalk helix splaying. (**C**) Importantly, the stalk helix backbone of our newly determined *Lp*MIP structure (blue) can be unambiguously placed in the 2Fo–Fc electron density map, shown here as a light blue mesh at 3σ. For clarity, only the density map for the stalk helix backbone is shown. (**D**) Overlay of the FKBP-like domains from the two *Lp*MIP structures. Residues surrounding the active site are shown as sticks, β-strands are labeled.

The splaying of the stalk helix, which emanates from the mid-helix residues ^76^EFNKK^80^, results in a relative reorientation of the attached FKBP-like domains in the two crystal structures. Nonetheless, both globular domains align with an RMSD of 0.214 Å (Fig. 1D). The main structural differences between the two FKBP-like domain structures were observed in the loop between β-strand 4 and 5, resulting in a different side-chain orientation for residue S189. Minor side-chain rearrangements were also seen for residues D142, V158 and Y185 in the active site which may however result from the different resolutions of the two structures.

Although microbial MIP proteins are promising drug targets, the structural similarity to human FKBP proteins raises concerns about possible cross-reactivity and off-target effects [34,35]. Naturally occurring inhibitors such as rapamycin (sirolimus) are large and chemically complex, poorly soluble in water, and have severe immunosuppressive effects limiting their use to treat microbial infection [36]. The comparison of human FKBP and pathogenic microbial MIP proteins bound to a chemically simpler, synthetic inhibitor molecules could thus present an important step towards improving ligand selectivity. Recently, an inhibitory effect of [4.3.1] bicylic sulfonamides on *L. pneumophila* proliferation in macrophages was demonstrated [34]. One such molecule, (1S,5S,6R)-10-((3,5- dichlorophenyl)sulfonyl)-5-(hydroxymethyl)-3-(pyridin-2-ylmethyl)-3,10-diazabicyclo [4.3.1]decan-2- one (JK095, Scheme 1), was co-crystallized with a human FKBP51 domain construct [34]. We thus deemed this compound a promising candidate for structural studies with MIP proteins from human pathogens and downstream structural comparison with human FKBPs. Isothermal titration calorimetry (ITC) confirmed that JK095 indeed interacts with microbial MIP proteins and *Lp*MIP variants (see below) and binds to full-length *Lp*MIP with a dissociation constant of 1.27 ± 0.14 µM (Fig. S1).

We also determined the structure of full-length *Lp*MIP in complex with JK095 by X-ray crystallography at 2.4 Å resolution (PDB: 8BJD) (Fig. 2A). The most notable structural differences between the crystal structures of *apo* and JK095-bound *Lp*MIP is the rearrangement of the loop connecting β-strands β4 and β5 near the stalk helix. Ligand binding to *Lp*MIP in solution was probed by titrating ^2^H, ^15^N-labeled *Lp*MIP with JK095 (Fig. 2B, C). Chemical shift perturbations were observed in the FKBP-like domain, consistent with the binding site identified in the crystal structure. In addition, residues within the FKBP domain facing the stalk helix, the stalk helix and the dimerization domain show chemical shift perturbations upon JK095 binding. The amide resonances between residues ∼57-76 in the N-terminal half of the *Lp*MIP stalk helix show severe line broadening and were thus not visible in the protein’s ^1^H, ^15^N-HSQC NMR spectrum (Fig. 2C, Fig. S2A). This suggests motions in the µs-ms timescale in this region. The FKBP-like domain shows complex shift changes upon JK095 addition, with some regions showing line broadening and others line sharpening. While crystallographic B-factors are generally less well suited to assess dynamic changes, overall, the changes in the presence of JK095 agree with the observed chemical perturbations in the NMR titrations. While this analysis is limited since the resolution of the two structures is incomparable, focusing on the changes of the distribution of individual B-values within individual structures together with the NMR data suggest dynamic quenching by the ligand throughout the protein (Fig. 2D, E).

**Fig. 2:**
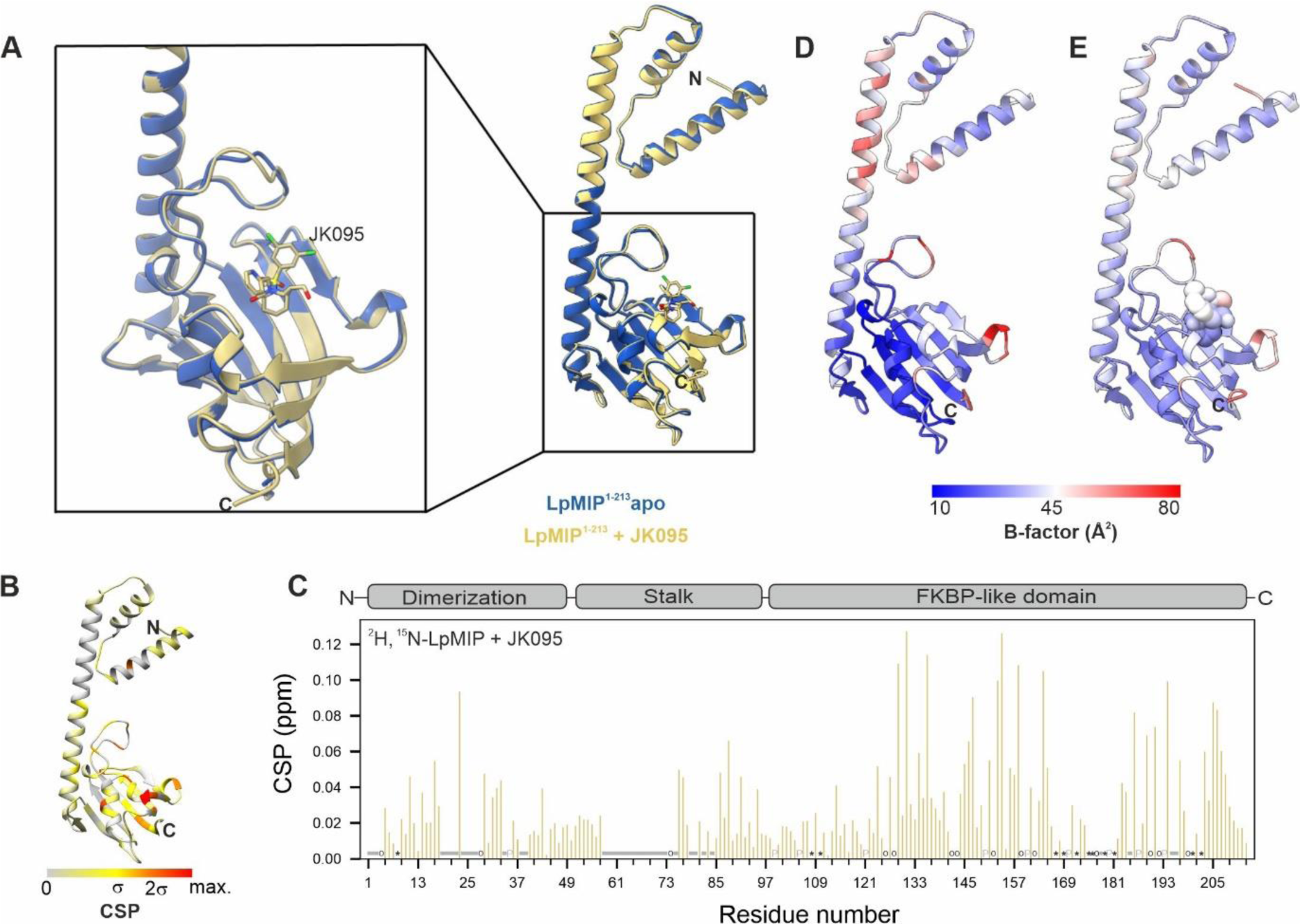
Comparison of full-length *Lp*MIP in the absence and presence of a bicyclic inhibitor. (**A**) Overlay of *Lp*MIP in the absence (blue, PDB: 8BJC) and presence of JK095 (yellow, PDB: 8BJD). The two structures align with a backbone RMSD of 0.349 Å. In the zoom of the FKBP-like domain, JK095 is shown as sticks. Non-carbon atom color scheme: blue: N, red: O, yellow: S, green: Cl. Note that the orientation of the zoom has been slightly tilted to better visualize the structural differences in the β4/β5-loop. (**B, C**) Chemical shift changes in ^2^H, ^15^N-labeled *Lp*MIP titrated with JK095 mapped on the *Lp*MIP crystal structure (B) and per residue (C) with the protein topology shown on top for orientation. Proline residues and residues without assignment in either state are labeled with grey P or indicated by a grey bar, respectively. Black circles (apo) and asterisk (JK095) represent resonances present only in one state. (**D, E**) Crystallographic B-factors of *Lp*MIP in the absence (D) and presence (E) of JK095.

To assess the structural dynamics of *Lp*MIP both locally and on a global scale in solution, we combined NMR relaxation studies with pulsed electron paramagnetic resonance (EPR) spectroscopy and small angle X-ray scattering (SAXS) (Fig. 3, Fig. S3-S6). NMR relaxation experiments informing on fast, ps-ns amide bond fluctuations and dynamics overlying the protein’s global rotational dynamics show that *Lp*MIP is relatively rigid on the assessed timescale, except for the very N-terminus, the linker between β3a and β3b, the linker between β4 and β5 and the C-terminus (Fig. S3). In contrast to the influence of JK095 on the protein dynamics on slower timescales, as was apparent through the changes in line broadening, fast backbone dynamics were not, or only marginally affected by the inhibitor.

**Fig. 3:**
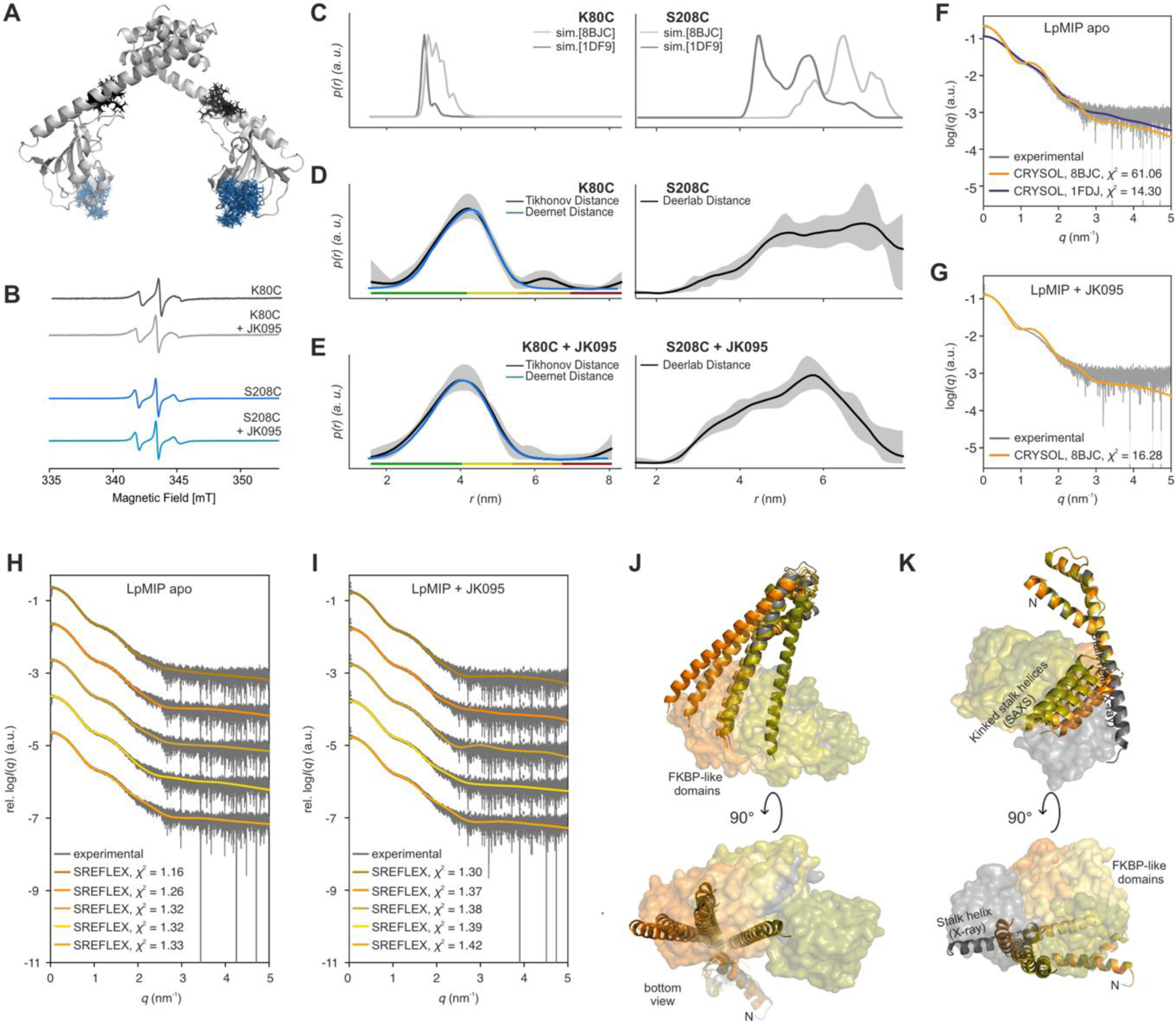
Structural dynamics of full-length *Lp*MIP in solution. (**A**) Simulated rotamers of proxyl-spin labels attached to *Lp*MIP at position K80C (black) or S208C (teal) (on PDB: 8BJC using MATLAB-based MMM2022.2 software). (**B**) Continuous-wave EPR spectra of spin-labeled *Lp*MIP single-cysteine variants. (**C**) Predicted interspin distances (sim.) for *Lp*MIP K80C (left) and *Lp*MIP S208C (right) based on the available *apo* state crystal structures (PDB-IDs: 8BJC, 1FD9). (**D, E**) Measured spin label distances using PELDOR/DEER spectroscopy in the absence (D) and presence (E) of JK095. For *Lp*MIP S208C, distances were determined through a global analysis of 4-pulse and 5-pulse PELDOR data (see Fig. S5). The rainbow code at the bottom indicates reliability for the probability distribution. (Green: shape, width and mean reliable; yellow: width and mean reliable, orange: mean reliable; red: not reliable) (**F, G**) SAXS scattering data for *Lp*MIP in the absence (F) and presence of JK095 (G). The simulated scattering curves (orange and blue traces) based on the available X-ray structures of *apo Lp*MIP (PDB: 8BJC, 1F9J) and with JK095 (PDB: 8BJD) do not match the scattering profile of the protein in solution after least-square fit to experimental values for 0.5 nm^-1^< *q* <1.5 nm^-1^. (**H, I, J, K)** For a better fit with the experimental SAXS data of *Lp*MIP in solution in the apo (H) and the JK095 bound state (I), SREFLEX modeling was carried out and yielded the calculated scattering profiles shown in the log plots and *Lp*MIP structural models with straight (J) and kinked (K) stalk helices. Accordingly, also the relative orientation of the FKBP like domains (shown as transparent surfaces) changes dramatically. The X-ray structure (PBD: 8BJC) is shown in grey, representative SREFLEX models in orange hues. For better visualization, models with straight and kinked helices are shown in separate chains. There are no discernible differences between the *apo* and JK095-bound state in the *Lp*MIP SREFLEX models, thus only the *apo* models are shown (for details see main text).

The results from EPR spectroscopy and SAXS further provide evidence of the high flexibility of *Lp*MIP in solution (Fig. 3). *Lp*MIP does not contain native cysteine residues. Thus, single cysteine mutants in the middle of the stalk helix (*Lp*MIP K80C) and at the C-terminal end of the FKBP-like domain (*Lp*MIP S208C) were introduced and labeled with nitroxide spin labels (Fig. 3A, Fig. S4, S5). Continuous wave EPR confirmed a satisfactory labeling efficiency at both positions (Fig. 3B).

Pulsed EPR spectroscopy (pulsed electron-electron double resonance (PELDOR, also known as DEER)) was used to determine the distances between the two spin-labeled sites, and the measurements were compared to simulations of the spin pair distance distributions based on the available crystal structures (Fig. 3C-E, Table S2). The distance distributions obtained from spin labeled *Lp*MIP K80C and S208C were broader than expected from the crystal structures, indicating that these structures represent only a subset of conformers in solution. Upon addition of JK095, no significant changes were observed for *Lp*MIP K80C, while for S208C the overall distribution shifted towards shorter distances. This could be explained e.g., by structural changes of the two FKBP domains moving closer together. Of note, the related NMR data show that at a molar protein:inhibitor ratio of 1:3 (n/n), the complex is already fully saturated. The EPR measurements were carried out with a protein:inhibitor ratio of 1:5, indicating that even when fully occupied, the “closed” conformation is only transiently populated.

Extensive structural dynamics of *Lp*MIP in solution are also apparent from SEC-SAXS experiments (Fig. 3F-K, Fig. S6, Table S3). Here, the *Lp*MIP scattering profiles did not match a simulated scattering curve using the available crystal structure, again suggesting a more complex conformational ensemble in solution. For a better fit with the experimental SAXS data of *Lp*MIP in solution, SREFLEX modeling was carried out [37] and *Lp*MIP structural models with straight and kinked stalk helices were obtained (Fig. 3J, K). While there were no discernible differences between the *apo* and JK095-bound state in the *Lp*MIP SREFLEX models, which may reflect the loss of JK095 during size exclusion chromatography (see below), the SAXS data show high domain flexibility concurrent with the EPR experiments.

### The appendage domains influence LpMIP dynamics and stability

Due to their high expression yields and solubility, deletion rather than full-length constructs have frequently been used for structural studies of both MIP and FKBP inhibitor complexes [21,38]. However, this may not only inadequately reflect the complexity of the therapeutic target, but also compounds a lack of understanding how the appendage domains affect protein structural dynamics and inhibitor binding. This question is exacerbated by our observation that ligand binding to the FKBP-like domains is sensed throughout the entire protein (Fig. 2).

In combination with our structural and spectroscopic studies on full-length *Lp*MIP, the modular architecture of *Lp*MIP provides a unique opportunity to explore such questions through deletion mutants. To emulate the structural diversity of MIP proteins from other human-pathogenic microbes, we generated two shortened *Lp*MIP constructs, *Lp*MIP^77-213^ and *Lp*MIP^100-213^ (Fig. S7A). *Lp*MIP^77-213^, containing the FKBP-like domain and a bisected stalk helix thus resembling *T. cruzi* MIP [26], is the construct typically used in *in vitro* ligand binding studies [20,21,39]. *Lp*MIP^100-213^, which consists solely of the FKBP domain, resembles e.g. *B. pseudomallei* MIP [33]. Both *Lp*MIP^77-213^ and *Lp*MIP^100-213^ are monomeric and structurally intact as seen by size exclusion chromatography (SEC) and circular dichroism (CD) spectroscopy (Fig. S7B-D). In a fluorescence-based assay, we saw that the melting temperature (*T_m_*) depended greatly on the protein’s appendage domains. (Fig. 4A). With 51.4 ± 0.3 °C, the *T_m_* of *Lp*MIP^100-213^ was found to be ∼14°C below that of the slightly longer construct *Lp*MIP^77-213^ (64.6 ± 0.6 °C) and ∼9 °C lower than that of full-length *Lp*MIP (60.7 ± 0.3 °C) (Fig 4A top). In all three constructs, addition of JK095 led to an increase in the melting temperature commensurate with protein stabilization upon inhibitor binding (Fig. 4A bottom). However, this effect was less pronounced for *Lp*MIP^100-213^ (Δ*T_m_*_(JK095-*apo*)_ = +2.8 °C) compared to both longer constructs (Δ*T_m_*_(JK095-*apo*)_ = +3.8 °C. This may reflect the strongly reduced binding affinity of JK095 to *Lp*MIP^100-213^ (*K_d_* = 20.47 ± 4.48 µM) compared to *Lp*MIP^77-213^ (*K_d_* = 2.27 ± 0.01 µM) and full-length *Lp*MIP (*K_d_* = 1.27 ± 0.14 µM) (Fig. S1). The differences in *T_m_* and inhibitor binding affinity suggest that the appendage domains, in particular the part of the stalk helix directly preceding the FKBP domain, play an important role in protein stability and ligand binding.

**Fig. 4:**
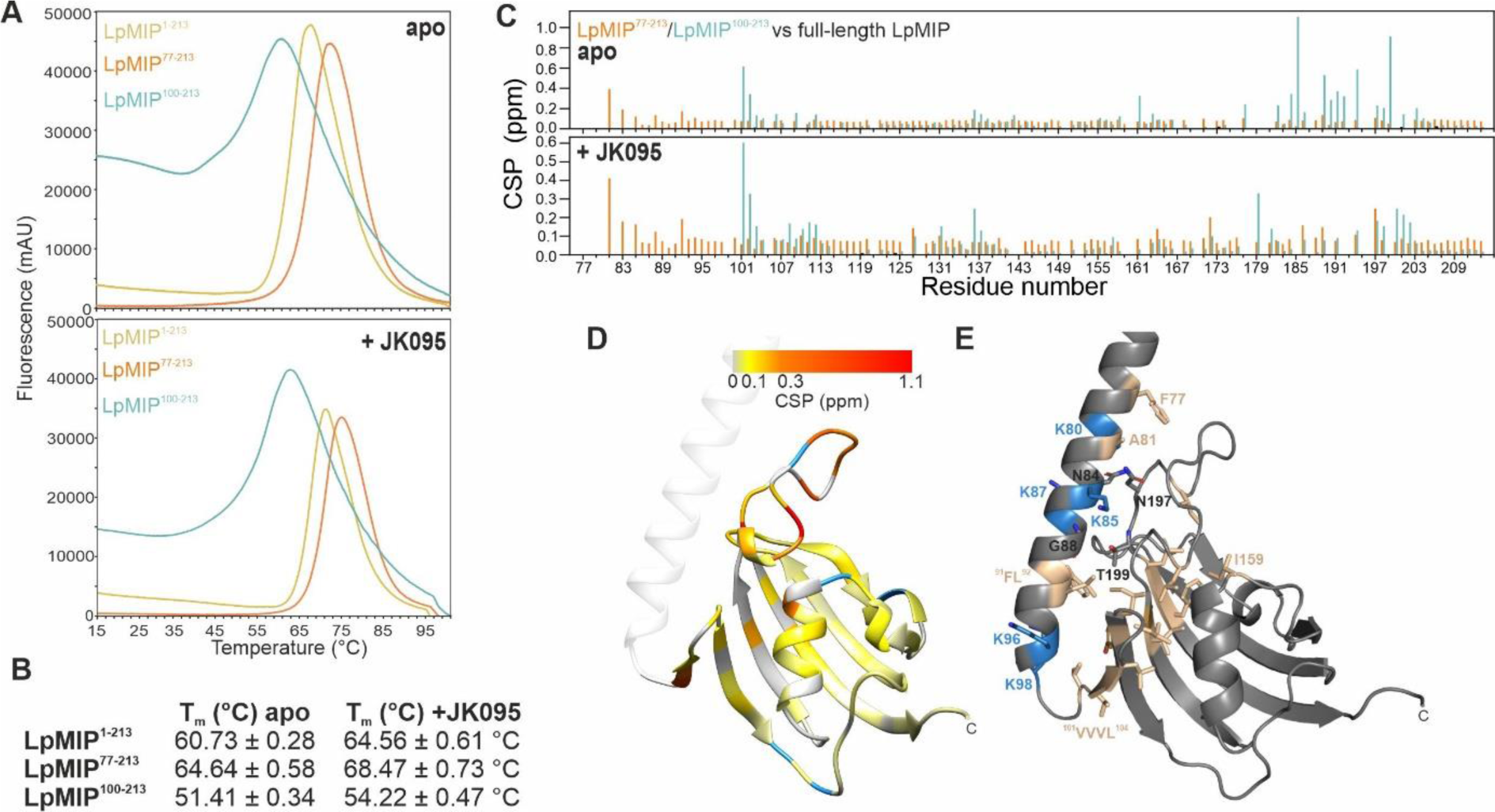
Role of the *Lp*MIP appendage domains for protein stability and crosstalk with the FKBP-like domain. (**A**) Fluorescence-based melting assay. The melting temperature (*T_m_*) for full-length *Lp*MIP (yellow) or two deletion constructs (orange, cyan) in the absence (top) or presence of a three-fold molar excess of JK095 (bottom) can be obtained from the inversion point of the upward slope. (**B**) *T_m_* values for the three constructs obtained from the curves shown in (A). Errors are standard deviations from three replicates. (**C**) Chemical shift perturbations of the FKBP-like domain resonances of *Lp*MIP^77-213^ and *Lp*MIP^100-213^ compared to full-length *Lp*MIP (orange and blue, respectively) in the *apo* state (top) and with JK095 (bottom). (**D**) Chemical shift differences between full-length *Lp*MIP and *Lp*MIP^100-213^ mapped on the FKBP-like domain, residues for which no signal is observed in *Lp*MIP^100-213^ are colored blue. (**E**) Details of hydrophobic interaction network between stalk helix and FKBP-like domain. Hydrophobic residues shown in sand, basic residues in blue, all others in grey. For a better overview, not all sidechains are shown.

To investigate the structural crosstalk between appendage and FKBP domains in *Lp*MIP in more detail, we used NMR spectroscopy. With the backbone assignments of all three *Lp*MIP constructs in the *apo* and JK095-bound states (Fig. S2), the chemical shifts for residues within the FKBP-like domains were compared (Fig. 4C, D). In the absence of inhibitor, there were only minor differences between full-length *Lp*MIP and *Lp*MIP^77-213^, except for the very N-terminal residues where the cleavage site is located (Fig 4C top, orange). Interestingly, differences between the two constructs became slightly more pronounced in the presence of JK095, particularly for residues 184 to 194 belonging to the β4/β5 loop (Fig 4C bottom, orange).

In contrast, the comparison between full-length *Lp*MIP with *Lp*MIP^100-213^ already showed strong chemical shift perturbations in the *apo* state (Fig 4C top, cyan). Most notable were the effects in the vicinity of residue 160 within the canonical ligand binding site, and between residues 180 and 200, which are part of the long loop between β-strands 4 and 5 and form an interaction network with the C-terminal half of the stalk helix (Fig. 4D, E). Furthermore, in the ^1^H, ^15^N-HSQC spectrum of *Lp*MIP^100-213^, no or extremely weak resonances for S115-N117, K146/T147, I159 and R188 were observed, while these were clearly visible in both longer constructs (Fig. 4D, E, Fig. S2). This suggests that these regions show altered dynamics in the absence of the stalk helix. However, except for residue I159 as well as R188 in the β4/5 loop, none of these residues are directly involved in FKBP/stalk helix interactions or part of the canonical ligand binding site, thus suggesting allosteric effects on the canonical binding site through the stalk helix. Potentially, such long-range crosstalk could be mediated through a hydrophobic interaction network between the stalk helix and FKBP-like domain (Fig. 4E).

Since the residues across all three full-length *Lp*MIP domains showed no significant differences in their respective backbone dynamics in the ps-ns timescale in {^1^H}^15^N-hetNOE experiments between the *apo* and the JK095-bound states (Fig. S3A), stalk helix removal seems to mostly affect slower, µs-ms motions within the FKBP-like domain. In the absence of the stalk helix, marginally increased hetNOE values for *Lp*MIP^77-213^ and *Lp*MIP^100-213^ could indicate slightly subdued backbone dynamics of the FKBP-like domain within the loops connecting β3a/β3b and β4/β5, both in the absence and presence of JK095 (Fig. S3).

### Role of the appendage domains for FKBP-like domain inhibitor binding

To gauge a possible structural role of the appendage domains for ligand binding in *Lp*MIP as suggested by our thermostability assays and NMR data (Fig. 4), we determined the crystal structures of *Lp*MIP^77-213^ (PDB: 8BK5) and *Lp*MIP^100-213^ (PDB: 8BK6) with JK095 at 2.26 and 1.49 Å resolution, respectively (Fig. 5A). These complement the crystal structure of full-length *Lp*MIP with JK095 (PDB: 8BJD, Fig. 2). The largest structural differences across all three *Lp*MIP constructs are observed in the β4/β5 loop, while the side chains of the active site residues adopted nearly identical orientations. JK095 bound to *Lp*MIP^77-213^ adopted a very similar binding stance as seen in the canonical binding pocket of full-length *Lp*MIP (Fig. 5A, B). However, in *Lp*MIP^77-213^, the inhibitor’s hydroxymethyl group adopted two orientations while in full-length *Lp*MIP, only the orientation facing away from the sidechain of D142 was observed, thereby forgoing the formation of a possible hydrogen bond interaction. Furthermore, the pyridine ring nitrogen was 2.7 Å away from the Y185 sidechain hydroxyl group in *Lp*MIP^77-213^, while this distance increased to 3.7 Å in full-length *Lp*MIP.

**Fig. 5:**
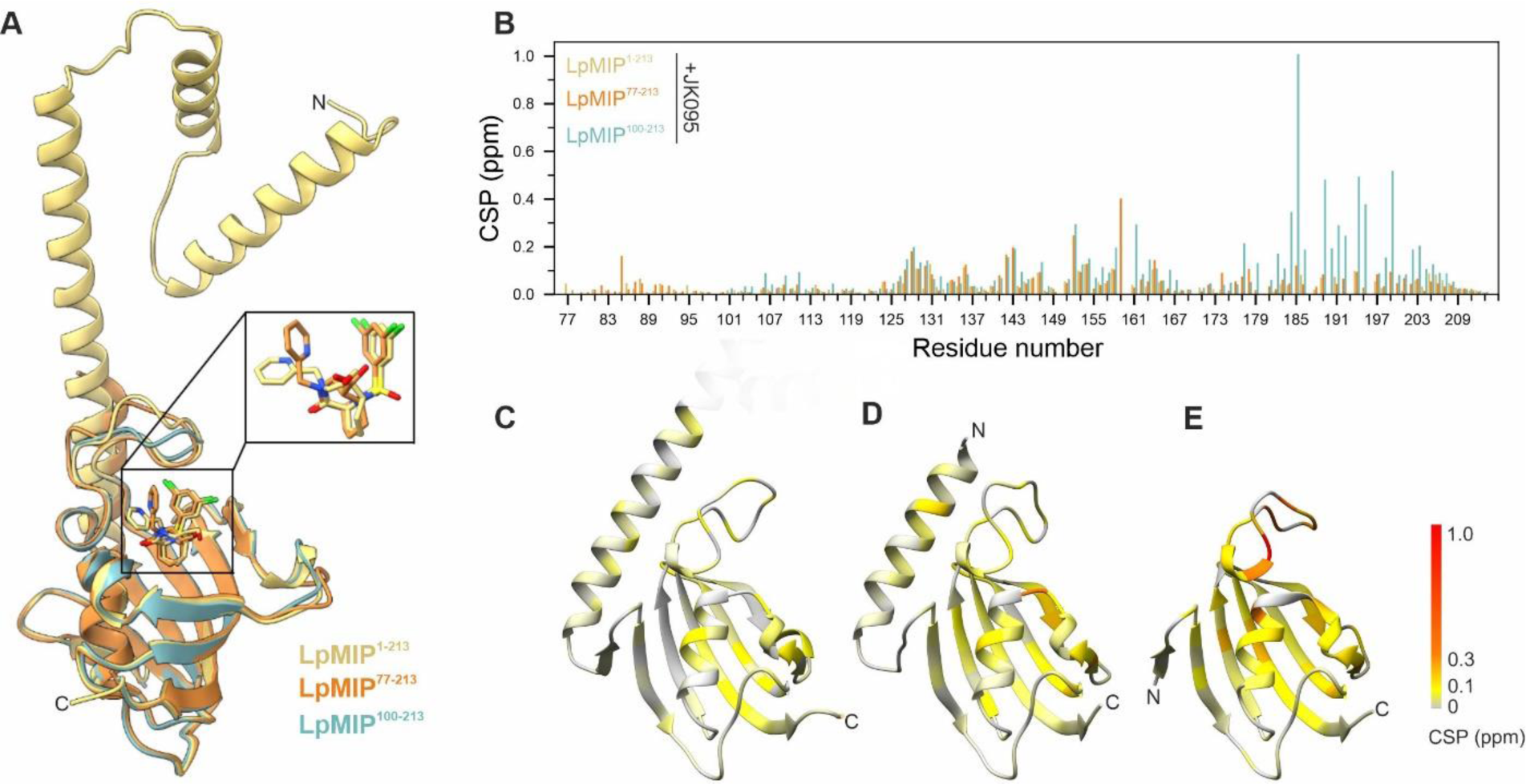
Stalk helix affects interaction of *Lp*MIP FKBP-like domain with a [4.3.1]-aza-bicyclic sulfonamide inhibitor. (A) Overlay of the X-ray crystal structures of *Lp*MIP^1-213^ (full-length), *Lp*MIP^77-213^ and *Lp*MIP^100-213^ co- crystallized with JK095 (PDB IDs: 8BJD, 8BK5, 8BK6). For the *Lp*MIP^1-213^ homodimer, only one subunit is shown. *Lp*MIP^100-231^ also crystallizes as a dimer, but no clear density for the ligand was obtained (for details see main text and compare Fig. S8). In the zoom-in, not that in *Lp*MIP^77-213^, the hydroxymethyl group of JK095 was found to adopt two different conformations. (B) Chemical shift perturbations in the FKBP-like domain of ^15^N-labeled full-length *Lp*MIP (yellow), *Lp*MIP^77-213^ (orange) and *Lp*MIP^100-213^ (teal) upon titration with JK095. For better comparison between the three constructs, a unified scale normalized to the maximal shift value in the FKBP-like domain across all three data sets was used. (**C-E**) JK095-induced chemical shift perturbations within the FKBP-like domain plotted on crystal structures of full-length *Lp*MIP (C), *Lp*MIP^77-213^ (D) and *Lp*MIP^100-213^ (E).

In contrast to the two longer constructs, the inhibitor binding site in *Lp*MIP^100-213^ was not clearly defined in the crystal structure (Fig. S8). To verify the possibility of drastically altered ligand interaction to the FKBP-like domain in the absence of the appendage domains in solution, we compared the chemical shift perturbations of the three ^15^N-labeled *Lp*MIP constructs titrated with JK095 (Fig. 5B-E, Fig. S2A-C). As expected, the chemical shift changes in full-length *Lp*MIP and *Lp*MIP^77-213^ agree with the binding site observed in the respective complex crystal structures. In stark contrast, addition of JK095 to *Lp*MIP^100-213^ affected a significantly larger number of residues and the chemical shift perturbation pattern was not restricted to the canonical ligand binding site. Of note, *Lp*MIP^100-213^ crystallized as a parallel dimer with the loop between β4 and β5 mediating many of the dimer contacts (PDB: 8BK6, Fig S8). These loops showed the largest structural differences between the two *Lp*MIP^100-213^ protomers in the unit cell and the largest chemical shift changes upon addition of JK095 in the NMR experiments. We thus wondered whether transient oligomerization could be responsible for the extensive JK095-dependent chemical shift perturbations in the 12 kDa *Lp*MIP^100-213^ construct. Under the assumption of isotropic tumbling, a rotation correlation time τ_c_ of 5.6 ns can be approximated according to the Stokes-Einstein equation for a spherical globular, monomeric protein of that size at 25 °C (see material and methods for details). By applying an empirical formula [40], a τ_c_ value of 7.3 ns can be derived for a 12 kDa molecule. Accordingly, neither the overall narrow line widths in the NMR spectra of ^15^N-labeled *Lp*MIP^100-213^ (Fig. S2C), nor the experimentally determined rotation correlation times (τ_c_ = 6.8 ± 0.9 ns for the *apo* protein, τ_c_ = 6.4 ± 0.7 ns in the presence of JK095) are indicative of inhibitor-induced dimer formation of *Lp*MIP^100-213^. Rather, the extensive NMR chemical shift perturbations in *Lp*MIP^100-213^ upon addition of JK095 are likely caused by the non-specific interaction with the inhibitor. This finding supports the notion that the *Lp*MIP appendage domains, particularly the C-terminal half of the stalk helix, play a decisive role in ligand binding to and dynamics within the FKBP domain.

### Comparison of *Lp*MIP and human FKBP51 in complex with the same [4.3.1]-aza-bicyclic sulfonamide inhibitor

*Lp*MIP^77-213^ shares 32 % sequence similarity with a construct of human FKBP51 (residues 16-140) that was recently co-crystallized with JK095 [41]. The two complex crystal structures (PDB IDs: 5OBK, 8BK5) align with a backbone RMSD of 0.776 Å (Fig. 6A). All residues interacting with JK095 are conserved between the two proteins (Fig. 6B). A conserved tyrosine residue (Y113/Y185 in FKBP51/*Lp*MIP) responsible for forming a H-bond to the nitrogen of the pyridine or bicycle of the inhibitor adopted the same orientation in both proteins. The sidechain of residue 159 forms a hydrophobic lid below the bi-cycle by forming van der Waals contacts with the inhibitor’s bi-cycle carboxy group. In addition, a barrage of aromatic residues in either protein nestles the bi-cyclic inhibitor core from below (Fig. 6B).

**Fig. 6:**
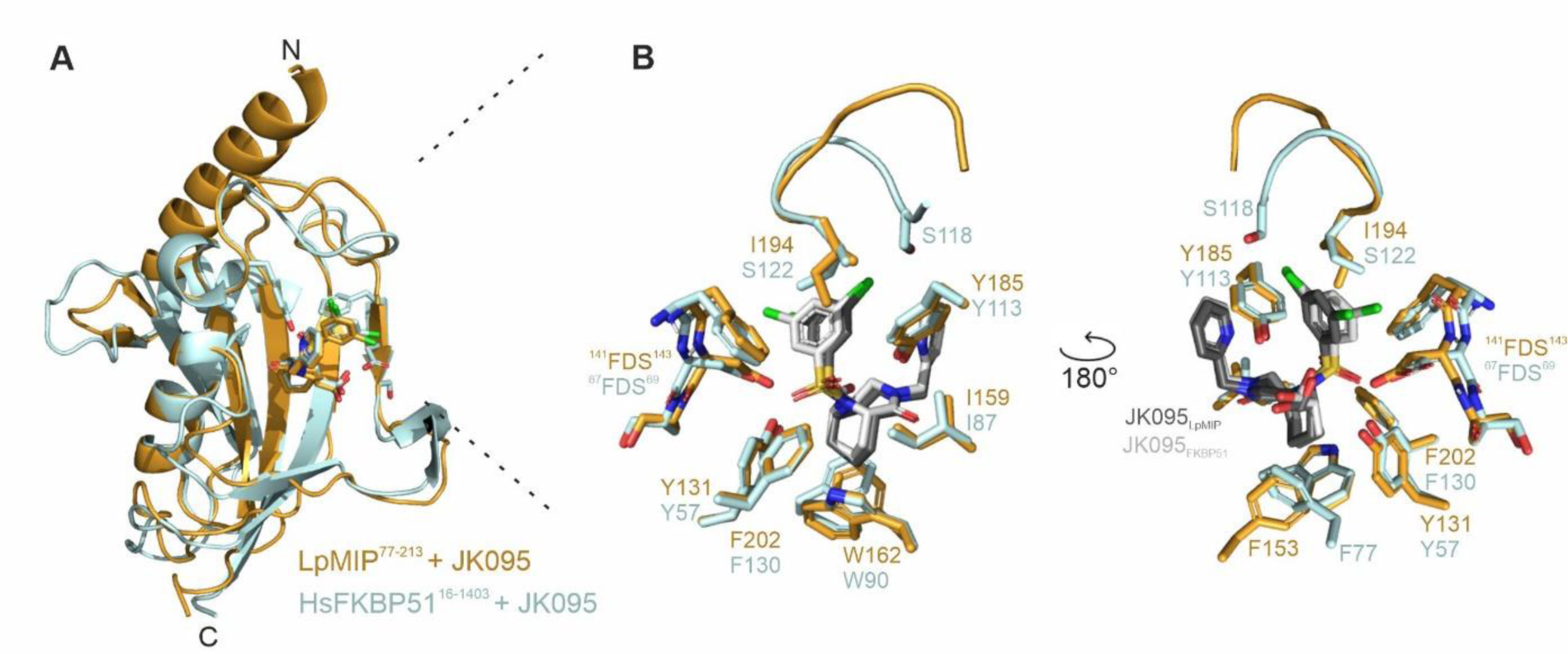
Comparison of *Lp*MIP and human FKBP51 in complex with the bicyclic inhibitor JK095. (A) Overlay of the crystal structures of *Lp*MIP^77-213^ (PDB: 8BK5, orange) and FKBP51^16-140^ (PDB: 5OBK, cyan) in complex with the [4.3.1]-aza-bicyclic sulfonamide JK095. (B) Zoom into the binding site. The relevant interacting residues are shown as sticks. JK095 is shown in dark (*Lp*MIP^77-213^) or light (FKBP51^16-140^) grey.

The inhibitor’s pyridine group, bi-cyclic core and sulfonamide group align well between the two proteins, only the di-chlorophenyl moiety is slightly differently tilted. Slight structural variations in the β3a-strand within the FKBP domain were found between FKBP51 and *Lp*MIP, namely across residues ^67^FDS^69^ and ^141^FDS^143^, respectively. The aromatic residue in this stretch may stabilize the di- chlorophenyl moiety through T-shaped π stacking. Inhibitor binding may also be affected by the structural and sequential differences in the loop connecting β4 and β5 (^117^GSLPKI^122^ in FKBP51 and ^189^SVGGPI^194^ in *Lp*MIP). Sitting on top of the di-chlorophenyl moiety of the ligand, the respective isoleucine residue within this stretch, together with the abovementioned phenylalanine in β3a, form a hydrophobic platform against which the di-chlorophenyl ring rests. In the case of FKBP51, the sidechain of S118 may additionally contact one chloro-substituent and thereby help to orient it. In contrast, the loop orientation observed in the *Lp*MIP^77-213^ crystal structure may disfavor interactions of either of the two chlorine groups with loop sidechains. The structural perturbation of the ^67/141^FDS^69/143^ motif in the β3a-strand also led to slightly different orientations of its central aspartic acid sidechain when comparing the structures of FKBP51^16-140^ and *Lp*MIP^77-213^. In both cases, the bound JK095 ligand’s hydroxymethyl group adopts two orientations. However, in FKBP51^16-140^, neither orientation comes close enough to form a hydrogen bond with the aspartic acid side chain of D68 (O-O distance 4.0 Å). In contrast, in *Lp*MIP^77-213^, in one of the two orientations the distance to the corresponding residue D142 is reduced by 0.9 Å compared to FKBP51^16-140^. In the other orientation, the inhibitor hydroxyl group can form hydrogen bonds with water molecules .

### Methylation leads to improved inhibitor binding to MIPs from different pathogenic microorganisms

It was recently observed that the stereospecific introduction of a methyl group at the C_α_ position of the pyridine substituent of bicyclic [4.3.1]-aza-amide inhibitors significantly increased their affinity for FKBP51 due to displacement of a surface water molecule [41]. JK095 does not carry such a methyl group and in our complex structure with *Lp*MIP^77-213^, we observed a crystallographic water in a similar surface position as the one that originally inspired the inhibitor methylation studies for human FKBP51 [41] (Fig. 7A). We thus wondered whether inhibitor methylation may be used to improve the affinity of bicyclic sulfonamides for MIP proteins from pathogenic microorganisms. To test this hypothesis, we introduced a methyl group into JK095, yielding JK236 (Scheme 1) and determined the co-crystal structure of *Lp*MIP^77-213^ with JK236 at 1.49 Å resolution (PDB: 8BJE) (Fig. 7B-D).

**Fig. 7:**
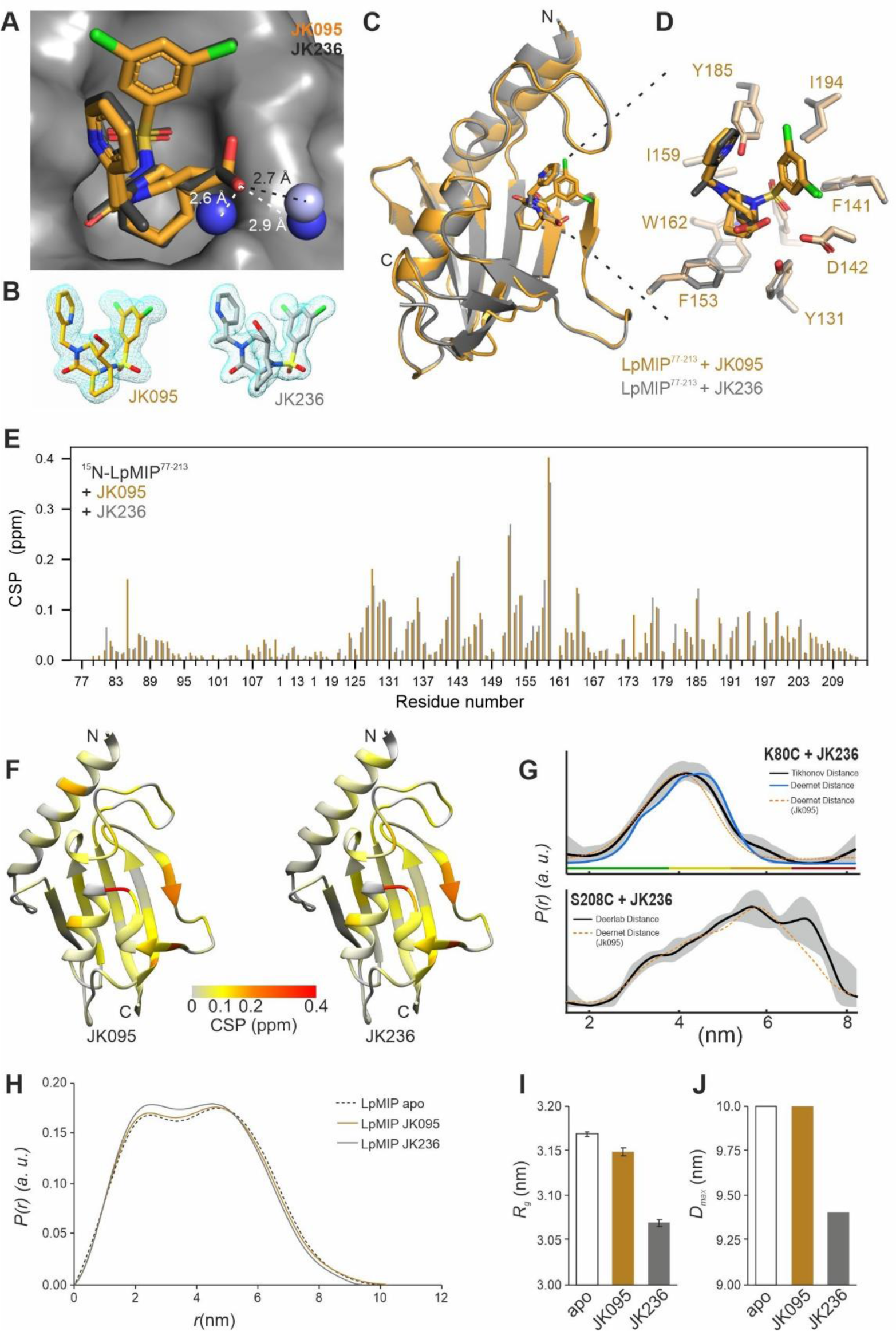
Solvent exposed methyl group in [4.3.1]-aza-bicyclic sulfonamide inhibitor improves affinity for *Lp*MIP^77-213^ through surface water displacement. (A) Water molecules in the crystal structures of *Lp*MIP^77-213^ with JK095 (PDB: 8BK5, dark blue spheres) and JK236 (PDB: 8BJE, light blue sphere). The additional methyl group in JK236 (pointing out of the paper plane) displaces one of the two water molecules that forms a hydrogen bond with the inhibitor’s hydroxymethyl group. Distances between crystallographic water and the inhibitors are indicated by white (JK095) and black (JK236) dashed lines. (B) Electron densities for the two inhibitor molecules in the co-crystal structures with *Lp*MIP^77-213^. Note that for JK095, the hydroxymethyl group adopts two conformations. (C) Overlay of the crystal structures of *Lp*MIP^77-213^ in complex with JK095 (PDB: 8BK5, orange) and its methylated derivative, JK236 (PDB: 8BJE, grey). For a structural comparison of the two molecules, see Scheme 1. (D) Zoom into the binding site. The relevant interacting residues are shown as sticks. (E) Relative NMR chemical shift perturbations (CSP) for JK095 (orange) and JK236 (grey) in comparison to the *apo* protein. (F) Chemical shift perturbation shown in (E) mapped on the X-ray structure of *Lp*MIP^77-213^ (PDB: 8BK5). (G) Measured spin label distances using PELDOR/DEER spectroscopy for spin-labeled full-length *Lp*MIP K80C (top) or S208C (bottom) with JK236. For better comparison, the distance distribution for JK095 (see Fig. 3) is indicated as a dashed orange line (without error margins). (H) SAXS derived real-space pair-distance distribution functions, or *p*(*r*) profiles, calculated for *Lp*MIP in the absence (dashed line) or presence of JK095 (orange line) or JK236 (grey line) and (**I, J**) resulting *R_g_* and *D_max_* values. *p*(*r*) functions were scaled to an area under the curve value of 1.

Overall, the structures of *Lp*MIP^77-213^ with JK095 and JK236 align with an RMSD of 0.283 Å and show no notable differences in protein sidechain or inhibitor conformations. Together with NMR chemical shift perturbation data of ^15^N-labeled *Lp*MIP^77-213^ titrated with JK095 or JK236 (Fig. 7E, F, Fig. S2D), this confirmed that both ligands interact in a highly similar fashion with the *Lp*MIP FKBP-like domain. Furthermore, pulsed EPR measurements of spin-labeled full-length *Lp*MIP K80C and *Lp*MIP S208C showed that JK236 affects the structural ensemble of full-length *Lp*MIP in a similar manner as JK095 (Fig. 7G, Fig. S5, S6).

Nonetheless, the binding affinity of JK236 to *Lp*MIP^77-213^ and full-length *Lp*MIP was increased by roughly one order of magnitude for the methylated (*K_d_* = 123.5 ± 47.4 nM and 108.5 ± 10.6 nM), compared to the unmethylated compound (2.27 ± 0.01 µM and 1.27 ± 0.14 µM) (Fig. S1). Presumably reflecting the increased affinity of JK236 over JK095, the SAXS data also show a more pronounced reduction in *R_g_* and *D_max_* for full-length LpMIP in the presence of the methylated inhibitor (Fig. 7H-J). Despite the presence of a less defined inhibitor interaction site in *Lp*MIP^100-213^, an increase in affinity was also observed or the shortest *Lp*MIP construct for the methylated ligand (*K_d_* = 20.47 ± 4.48 µM vs 1.31 ± 0.24 µM for JK095 and JK236, respectively).

A surface water molecule is indeed displaced in the JK236 co-crystal structure compared to the complex with JK095 (Fig. 7A). While the two inhibitors bound to *Lp*MIP superimpose nearly perfectly, the orientation of the hydroxymethyl group is fixed in JK236 in contrast to the two orientations observed for JK095. In JK236, the hydroxymethyl group faces away from the sidechain of D142 and instead exclusively forms a hydrogen bridge with a water molecule. At a resolution of 1.49 Å, the additional methyl group in JK236 can also be placed unambiguously in the crystal structure and is seen to point into the solvent where it does not undergo any protein contacts but rather displaces a water molecule (Fig. 7A). This shows that the methylation of bicyclic ligands to obtain high-affinity binders through surface water displacement is feasible for *Lp*MIP and may constitute a general concept for FKBPs as well as microbial MIPs.

To gauge whether methylation for improved binding is indeed applicable to MIPs from other human pathogens including those of eukaryotic origin, we turned to the protozoan *Trypanosoma cruzi*, the causative agent of Chagas disease. With a free-standing stalk helix and a prototypical FKBP domain, the *T. cruzi* MIP protein (*Tc*MIP) structurally resembles the *Lp*MIP^77-213^ construct lacking the dimerization domain and N-terminal half of the stalk helix (Fig. 8).

**Fig. 8:**
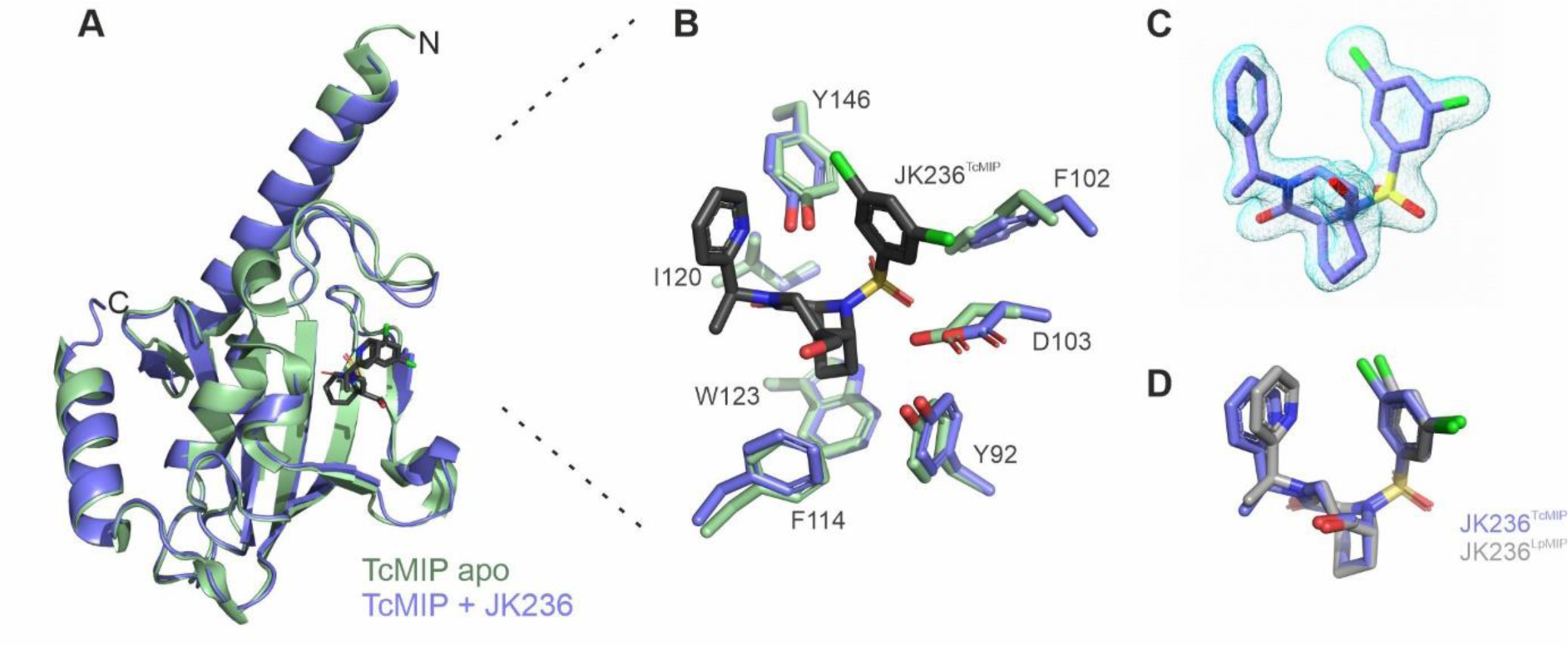
T*r*ypanosoma *cruzi* MIP in complex with a [4.3.1]-aza-bicyclic sulfonamide inhibitor. (A) Overlay of the crystal structures of *apo Tc*MIP (green, PDB: 1JVW) and JK236-bound *Tc*MIP (blue, PDB: 8BK4). (B) Active site residues in the *apo* or JK236-bound *Tc*MIP. The ligand is shown in black. (C) Electron density for JK236 bound to *Tc*MIP. The 2Fo-Fc electron density maps are shown in light blue mesh at 3σ. (D) Comparison of the inhibitor binding stance in *Tc*MIP (blue) and *Lp*MIP^77-213^ (grey). For details, see also Fig. S9.

Similar to *Lp*MIP, ligand binding to *Tc*MIP was improved for the methylated (*K_d_* = 45.5 ± 9.2 nM) versus the non-methylated compound (599.0 ± 25.5 nM) (Fig. S1). Our crystal structure of *Tc*MIP in complex with JK236 (PDB: 8BK4) at 1.34 Å resolution confirms the interaction of JK236 with the canonical binding site in the FKBP-like domain and a highly similar interaction mode as seen for *Lp*MIP (Fig. 8, Fig. S9).

The complex structure aligns to the previously published structure of *apo Tc*MIP (PDB: 1JVW) [26] with an RMSD of 0.499 Å (Fig. 8A). The largest differences between the two proteins are seen again in the loop connecting β-strands 4 and 5, as well as in β-strand 3a. In the *Tc*MIP *apo* structure, multiple water molecules are found around the substrate binding site which are absent with JK236, but no surface water molecule is seen in the same position as detected in JK095-bound FKBP51 [34] and *Lp*MIP. However, due to the lack of a complex structure of *Tc*MIP with JK095, it is difficult to assess the consequences of inhibitor methylation on water occupancy in *Tc*MIP in detail. Nonetheless, the similar gain in binding affinity through the introduction of the methyl group into the bi-cyclic inhibitor indicates a similar mode of action that can be exploited for the development of high-affinity binders against MIP proteins from various pathogens. The availability of two structures of MIP proteins from highly diverse pathogenic microorganisms in complex with the same synthetic inhibitor now also provides a unique opportunity to elucidate the possibility to generate pan-inhibitors.

## Discussion

The role of MIPs as widespread microbial virulence factors has spurred efforts to develop inhibitors targeting the MIP FKBP-like domain as the most conserved MIP domain. However, many MIP proteins contain additional appendage domains of unknown function. This prompted us to investigate the interdomain crosstalk and dynamics of the homodimeric *Legionella pneumophila* MIP protein as a representative model system for multi-domain MIPs in more detail.

Intrinsic structural flexibility seems to be a hallmark of homodimeric MIP proteins from pathogenic microorganisms [42]. Not only did we notice significant stalk helix splaying between the two available crystal structures of full-length *Lp*MIP in the absence of a ligand, but a recently published structure of unliganded, homodimeric *P. aeruginosa* FkbA, which shares the same three-domain architecture, showed both straight and bent stalk helices in the crystal structure [25]. It has been suggested that variations in crystal structures are a good proxy for dynamics in solution [43] and in the case of *Lp*MIP, we can support and extend this notion with EPR and NMR spectroscopy as well as SAXS. Our crystal structures provide a glimpse of the protein’s dynamics, but the full extent of its domain gymnastics in solution required a multi-faceted approach.

Using NMR spectroscopy, we identified a dynamic hotspot in the central stalk helix of *Lp*MIP. This is also the region that shows extensive kinking in our SAXS SREFLEX models. A difference in ”bending“ of the central stalk helix was mentioned previously for a co-crystal of full-length *Lp*MIP with FK506 [14], although the corresponding data set has never been submitted to the PDB and thus cannot be analyzed in detail here. Pervushin and colleagues reported that the *E. coli* FkpA stalk helix rigidifies in the presence of a client protein and led to reduced interdomain mobility [42]. Here, we saw that binding of a bi-cyclic vinylsulfone inhibitor led to complex changes throughout the protein, possibly including the rigidification of the N-terminal half of the stalk helix.

Comparing JK095-bound *Lp*MIP^77-213^ with the rapamycin-bound protein (Fig. S10), shows the relative displacement of the ligand enclosing sidechains and indicates that the active site of *Lp*MIP displays a conformational flexibility commensurate with its ability to bind to differently sized ligands. Across all our structures, the β4/β5 loop, which interacts with the stalk helix and may thus serve as a substrate-selective communication node between stalk and FKBP-like domain, showed the most structural variations. In contrast to previous observations with rapamycin [21], no significant rigidification of FKBP-like domain loops on very fast timescales was observed with JK095, while slower dynamics were quenched throughout the protein upon ligand binding. Different inhibitor molecules could thus potentially mimic the structural and dynamic consequences of diverse, yet unidentified, native ligands. Unfortunately, the affinity of collagen peptides, the only known native *Lp*MIP substrate to date [19,20], is too low for detailed structural and dynamic analysis.

Furthermore, the addition of bi-cyclic inhibitors led to a population shift but not a full transition to a “closed” conformation with decreased distances between the FKBP-like domains in our EPR experiments. Whether this is a general feature of *Lp*MIP ligands or unique to the tested inhibitors is unknown. Future ligand screening could explore whether the ability of ligands to shift the *Lp*MIP conformational ensemble to a closed state correlates with its antimicrobial efficiency.

We could also show that the *Lp*MIP domains engage in bidirectional crosstalk. Ligand binding at the FKBP-like domain affected the stalk helix and dimerization domain, and, in turn, stalk helix deletion reduced protein stability and, surprisingly, led to the loss of a defined ligand binding mode. The allosteric modulation of ligand binding by the C-terminal half of the stalk helix has interesting implications for ligand recognition and regulation of MIP proteins from other pathogenic species, such as *Burkholderia pseudomallei*, which naturally lack a stalk helix and dimerization domain [33].

Deletion constructs of MIP proteins have been commonly used to study inhibitor binding. Our data suggests that a construct retaining the C-terminal half of the stalk helix is suitable for most applications, but there are nonetheless some differences to consider. The increased melting temperature of *Lp*MIP^77-213^ may indicate that stabilization of the FKBP domain by the stalk helix’ C-terminal end is counteracted by the protein’s flexibility in the N-terminal half. Complete deletion of the stalk helix has negative consequences for both protein stability and ligand interactions.

Bi-cyclic sulfonamides have antiproliferative effects against *L. pneumophila* and *Chlamydia pneumoniae*, which both express MIP proteins [34]. This suggests that the bicyclic sulfonamide scaffold is a promising starting point for drug development. Our results on *T. cruzi* MIP suggest that both prokaryotic and eukaryotic MIP proteins can be targeted with a high-affinity pan-inhibitor, and lessons from human FKBPs such as site-specific methylation [41] can be exploited to improve inhibitor affinity for microbial MIPs. However, the structural similarities between MIPs and FKBPs pose challenges, particularly since FKBP inhibition leads to immunosuppression, the opposite of the desired effect in fighting severe infections. Here, we could carry out a structural comparison of a microbial MIP with a human FKBP in complex with the same synthetic ligand for the first time. In a previous NMR study on FKBP51, the central aromatic residue in the β3a-strand, was seen to flip in and out of the binding pocket, a process important for ligand selectivity [44]. The residues stabilizing the “outward” position (FKBP51 K58, K60 and F129) are not fully conserved in *Lp*MIP (T132, R134, F202). Hence ring flipping might be an important distinguishing feature between the two proteins. Additional structures and dynamic studies of human FKBPs and microbial MIPs in complex with the same ligands, possibly with other molecular scaffold architectures, may be helpful in making further progress in this area.

In summary, we found that in *Legionella pneumophila* MIP, the stalk helix decisively modulates ligand-binding behavior of the FKBP-like domain, the most conserved domain across all MIP proteins. This, together with the high intrinsic flexibility of MIP proteins and the ability to engage with structurally diverse ligands, suggests that MIP appendage domains can be used to fine-tune substrate responses and suggest they play a contextual role in the survival and replication of pathogenic microorganisms.

## Material and Methods

### Cloning, protein expression and purification

Genes coding for *Legionella pneumophila Lp*MIP^1-213^, *Lp*MIP^77-213^, *Lp*MIP^100-213^ and *Trypanosoma cruzi Tc*MIP with a His_6_-tag were obtained from GenScript (Piscataway Township, NJ, USA) and cloned into a pET11a vector. Single cysteine mutants for EPR spectroscopy were introduced at positions K80C and S208C in *Lp*MIP^1-213^ via site directed mutagenesis using the following primer pairs: K80C forward: 5’-CCGCGGAGTTTAACAAGTGCGCGGATGAAAACAAGG-3’ K80C reverse 5’- ACCTTGTTTTCATCCGCGCACTTGTTAAACTCCGCG–3’ S208C forward 5’- TAAGATTCACCTGATCTGCGTGAAGAAAAGCAG – 3’ S208C reverse 5’- CTGCTTTTCTTCACGCAGATCAGGTGAATCTTA – 3 Freshly transformed *E coli.* BL21 gold (DE3) cells were grown at 37 °C to an OD_600_ of 0.6 and then induced with 1 mM IPTG and grown overnight at 20 °C. ^2^H, ^15^N-labeled *Lp*MIP^1-213^ was obtained by growing cells in commercially available Silantes OD2 *E. coli* medium (Silantes GmbH, Munich, Germany). ^13^C, ^15^N-labeled *Lp*MIP^77-213^ and *Lp*MIP^100-213^ were obtained by growing cells in minimal medium with ^15^N-NH_4_Cl and ^13^C-glucose as the sole nitrogen and carbon sources. Cells were harvested by centrifugation (5000×g, 10 min, 4 °C). The cell pellet was frozen in liquid nitrogen and stored at – 20 °C until further use.

For purification of *Lp*MIP^1-213^ and *Lp*MIP^77-213^, the cell pellet was dissolved in lysis buffer (20 mM Tris pH 8, 20 mM Imidazole pH 8, 300 mM NaCl, 0.1 % Tx100, 1 mM DTT, 1 mM benzamidine, 1 mM PMSF, DNAse, RNAse and lysozyme). Cells were disrupted passing them three times through a microfludizer (Maximator) at 18,000 psi. Membranes and cell debris were pelleted at 48,380xg, 30 min, 4 °C and the supernatant was loaded onto a NiNTA column (Qiagen, Hilden, Germany) previously equilibrated with washing buffer (20 mM Tris pH 8, 300 mM NaCl and 20 mM imidazole). After washing with 10 CV (column volumes) of washing buffer, the protein of interest was eluted with 5 CV of elution buffer (20 mM Tris pH 8, 300 mM NaCl and 500 mM imidazole pH 8). Proteins were dialyzed overnight at 4 °C in 20 mM Tris pH 8, 300 mM NaCl in the presence of His-tagged TEV protease (1:20 mol/mol) to cleave the His-tag from the MIP constructs. Dialyzed protein was then loaded onto a fresh NiNTA column. The flow through was collected and the column was washed with 4 CV of washing buffer to obtain the maximum amount of tag-free MIP protein. For the purification of *Lp*MIP^100-213^ the same protocol was applied, with all buffers adjusted to pH 7. After concentration, the proteins were loaded on a size exclusion column (HiLoad 16/600 Superdex 200 pg, Cytiva, Freiburg, Germany) equilibrated with size exclusion buffer (20 mM Tris pH 7, 150 mM NaCl for *Lp*MIP^77-213^ and *Lp*MIP^100-213^ and 50 mM Tris pH 7, 150 mM NaCl for *Lp*MIP^1-213^). The fractions containing pure protein were pooled and sample purity was verified by SDS-PAGE.

### Crystallization, data collection and structure determination of *Lp*MIP inhibitor complexes

Following size exclusion chromatography, each of the proteins were kept in a solution of 20 mM Tris and 150 mM NaCl at pH 7.0 and were concentrated to 10 mg/mL using a 10,000 MWCO concentrator. Each protein was mixed with the crystallization buffer in a ratio of 2:1, and, where appropriate, with a 1:5 molar ratio of inhibitor. Inhibitors were synthesized as previously described [34,41]. All crystals were obtained using sitting drop v*apo*r diffusion via custom screens with the following conditions: *Lp*MIP^1-213^ 20 % (w/v) PEG 6000, 500 mM zinc acetate dihydrate, 100 mM MES, pH 6.0. *Lp*MIP^1-213^JK095 15 % (w/v) PEG 6000, 500 mM zinc acetate dihydrate, 100 mM MES, pH 6.5. *Lp*MIP^100-213^ JK095 20 % (w/v) PEG 8000, 500 mM zinc acetate dihydrate, 100 mM MES, pH 5.8. *Lp*MIP^77-213^ JK095 20 % (v/v) 2-propanol, 0.2 M sodium citrate tribasic dihydrate, 0.1 M HEPES, pH 7.5. *Lp*MIP^77-213^ JK236 18 % (w/v) PEG 8000, 0.2 M zinc acetate, 0.1 M sodium cacodylate, pH 6.5. *Tc*MIP JK236 30 % (v/v) MPD, 0.2 M ammonium acetate, 0.1 M sodium citrate, pH 5.6. Crystals were briefly soaked in 30 % (v/v) glycerol for cryo-protection and subsequently flash-frozen in liquid nitrogen in preparation for diffraction experiments at synchrotron energy. Data were collected at beam line ID23-1 and ID30A- 3 (ESRF, Grenoble).

Crystals of the MIP series diffracted between 1.3 and 2.4 Å resolution (Table 1). Data were processed with XDS [45] and structures were solved by Molecular Replacement with Phaser [46] using previously published models of MIPs (PDB ID: 1FD9, 1JVW). Manual rebuilding was performed with COOT [47] and refinement with Refmac [48]. The refined models were deposited into the PDB repository with the following IDs: 8BJC, 8BJD, 8BJE, 8BK4, 8BK5, 8BK6. Images were prepared using Pymol (Schrödinger, LLC), CorelDRAW (Corel), UCSF ChimeraX [49] and Blender (Blender Foundation).

### Analytical Size-exclusion chromatography (SEC)

20 µM of purified *Lp*MIP constructs (*Lp*MIP^1-213^, *Lp*MIP^77-213^ or *Lp*MIP^100-213^) in 20 mM Tris pH 7, 150 mM NaCl were used. For the *apo* state protein, a final concentration 0.02 % DMSO was added. A 5-fold molar excess of JK095 in DMSO was added (0.02 % final DMSO concentration). Samples were injected on a Superdex200 Increase 10/300 GL (Cytiva) column via an NGC chromatography system (BioRad).

### Circular Dichroism (CD) spectroscopy

CD measurements were conducted on a Jasco J-1500 CD spectrometer (Jasco, Gross-Umstadt, Germany) with 1 mm quartz cuvettes using 3.5 µM protein in 5 mM Tris pH 7 and 2.5 mM NaCl. Spectra were recorded at 25 °C in a spectral range between 190 – 260 nm with 1 nm scanning intervals, 1.00 nm bandwidth and 50 nm/min scanning speed. All spectra were obtained from the automatic averaging of five measurements.

### Isothermal Titration Calorimetry (ITC)

Experiments were performed in an isothermal titration calorimeter (Microcal ITC200 - Malvern Panalytical) at 25 °C with a reference power of 11 µCal/sec, an initial delay of 120 seconds and a stirring speed of 750 rpm. Protein concentration within the cell was between 20 and 40 µM and ligand concentration in the syringe was between 0.5 and 1 mM. Protein and inhibitors (JK095 and JK236) were prepared in 20 mM Trips pH 8, NaCl 150 mM and 5 % DMSO. For each titration, 20 injections (spacing between injections was 180 sec, duration was 0.4 sec) of 2 µL inhibitor solution were carried out. The curves were fitted using Origin.

### NMR spectroscopy

All NMR spectra were obtained at 298.2 K on 600 MHz Bruker AvanceIII HD or Neo NMR spectrometer systems equipped with 5-mm triple resonance cryo-probes. The spectrometers were locked on D_2_O. The ^1^H chemical shifts of the ^2^H, ^15^N-labelled *Lp*MIP^1-213^, ^13^C, ^15^N-labelled *Lp*MIP^77-213^ and ^13^C, ^15^N-labelled *Lp*MIP^100-213^ were directly referenced to 3-(trimethylsilyl)propane-1-sulfonate (DSS). ^13^C and ^15^N chemical shifts were referenced indirectly to the ^1^H DSS standard by the magnetogyric ratio [50]. *Lp*MIP^1-213^ was measured in 50 mM Tris HCl pH 7, 150 mM NaCl, 0.1 mM DSS, 0.05 % NaN_3_ and 10 % D_2_O. Sample conditions for *Lp*MIP^77-213^ and *Lp*MIP^100-213^ were the same except 20 mM Tris HCl pH 7 was used. Final protein concentrations were in the range of 100-150 µM. All spectra were processed using Bruker Topspin 4.1.1 and analyzed using CcpNmr Analysis [51] v2.5 (within the NMRbox virtual environment [52]).

The previously published NMR backbone assignments of *Lp*MIP^1-213^ (BMRB entry 7021) and *Lp*MIP^77-213^ (BMRB entry 6334)^37,38^ were transferred to our spectra and verified using band-selective excitation short-transient (BEST) transverse relaxation-optimized spectroscopy (TROSY)-based HNCA or HNCACB experiments under our buffer conditions. In contrast, the assignment of *Lp*MIP^100-213^ had to be determined *de novo* by a set of BEST-TROSY-based HN(CA)CO, HNCA and HN(CO)CA, as the ^1^H, ^15^N-HSCQ spectrum of this construct differed significantly from the resonances of the FKBP domain in both *Lp*MIP^77-213^ and full-length *Lp*MIP.

Standard NMR pulse sequences implemented in Bruker Topspin library were employed to obtain *R_1_*, *R_2_* and *^15^N,{^1^H}-NOE* values. For *Lp*MIP^1-213^, TROSY-sampling pulse sequences were used to ensure high data quality. Longitudinal and transverse ^15^N relaxation rates (*R_1_* and *R_2_*) of the ^15^N-^1^H bond vectors of backbone amide groups were extracted from signal intensities (*I*) by a single exponential fit according to equation **1**:

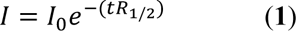

In *R_1_* relaxation experiments the variable relaxation delay *t* was set to 1000 ms, 20 ms, 1500 ms, 60 ms, 3000 ms, 100 ms, 800 ms, 200 ms, 40 ms, 400 ms, 80 ms and 600 ms. In all *R_2_* relaxation experiments the variable loop count was set to 36, 15, 2, 12, 4, 22, 8, 28, 6, 10, 1 and 18. The length of one loop count was 16.96 ms. In the TROSY-based *R_2_* experiments the loop count length was 8.48 ms. The variable relaxation delay *t* in *R_2_* experiments is calculated by length of one loop count times the number of loop counts. The inter-scan delay for the *R_1_* and *R_2_* experiments was set to 4 s.

The ^15^N-{^1^H} steady-state nuclear Overhauser effect measurements (*^15^N,{^1^H}-NOE*) were obtained from separate 2D ^1^H-^15^N spectra acquired with and without continuous ^1^H saturation, respectively. The *^15^N,{^1^H}-NOE* values were determined by taking the ratio of peak volumes from the two spectra, *^15^N,{^1^H}-NOE* = *I_sat_/I_0_*, where *I_sat_* and *I_0_* are the peak intensities with and without ^1^H saturation. The saturation period was approximately 5/*R_1_* of the amide protons.

The averaged ^1^H and ^15^N weighted chemical shift perturbations (CSP) observed in ^1^H, ^15^N-HSQC spectra were calculated according to equation **2** [53]:

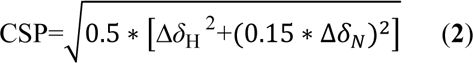

Here, ΔδH is the ^1^H chemical shift difference, ΔδN is the ^15^N chemical shift difference, and CSP is the averaged ^1^H and ^15^N weighted chemical shift difference in ppm.

The oligomerization state of a protein can be estimated from the rotational correlation time (τ_*c*_), the time it takes the protein to rotate by one radian under Brownian rotation diffusion. Under the assumption of a spherical globular protein and isotropic motion, τ_*c*_ (in ns) can be roughly approximated from the Stokes-Einstein equation (**3**):

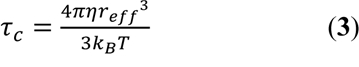

where η is viscosity (0.89 mPa*s for water at 298.2 K), *k_B_* the Boltzmann constant and *T* the absolute temperature. The effective hydrodynamic radius *r_eff_* can directly be correlated with molecular weight (*M_w_*):

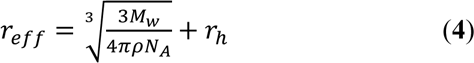

where ρ is the average protein density (1.37 g/cm^3^) and *N_A_* the Avogadro constant. For our calculations we used hydration layer radius of 3.2 Å.

Based on studies from the Northeast Structural Genomics Consortium an empirical formula could be derived for direct correlation of *M_w_* (in Da) and τ_*c*_ (in ns) for proteins in the range of 5-25 kD [40]:

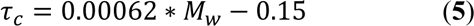

The rotational correlation time is directly accessible from the ratio of ^15^N *R_1_* and *R_2_* relaxation rates of backbone amide measured at a ^15^N resonance frequency (*ν*_*N*_) assuming slow isotropic overall motion [40,54] (equation **6**):

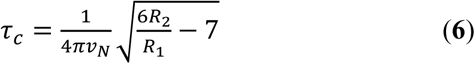

### Electron Paramagnetic Resonance (EPR) spectroscopy sample preparation

For spin labelling, Ni-NTA-column-bound single cysteine mutants of *Lp*MIP^1-213^ were incubated overnight at 4 °C using a 15-fold excess of 3-(2-Iodoacetamido)-2,2,5,5-tetramethyl-1-pyrrolidinyloxy (IPSL) after the washing steps and then purified as described above. Following the IPSL-labelling procedure, 4 µL of D_8_-glycerol or water was added to a 12 µL of *Lp*MIP sample, mixed thoroughly and gently transferred into a sample tube. The samples for continuous wave EPR were directly measured in a 25 μL micropipettes (BRAND, Germany) with a 0.64-mm diameter at room temperature. Samples for pulsed EPR were flash frozen in liquid nitrogen in a 1.6 mm quartz EPR tube (Suprasil, Wilmad LabGlass) and stored at -80°C.

### Continuous-wave EPR measurements

Continuous-wave (CW) EPR measurements were performed at X-band frequency (9.4 GHz) on a Bruker EMXnano Benchtop Spectrometer at room temperature in a 25 µL micropipette (BRAND, Germany) with a 0.64 mm diameter. The spectra were acquired with 100 kHz modulation frequency, 0.15 mT modulation amplitude, 0.6 - 2 mW microwave power, 5.12 ms time constant, 22.5 ms conversion time, and 18 mT sweep width.

### Pulsed EPR measurements

Pulsed EPR (PELDOR/DEER) experiments were performed on a Bruker Elexsys E580 Q-Band (33.7 GHz) Pulsed ESR spectrometer equipped with an arbitrary waveform generator (SpinJet AWG, Bruker), a 50 W solid state amplifier, a continuous-flow helium cryostat, and a temperature control system (Oxford Instruments). Measurements were performed at 50 K using a 10 – 20 µL frozen sample containing 15 – 20 % glycerol-*d_8_* in a 1.6 mm quartz ESR tubes (Suprasil, Wilmad LabGlass). For measuring the phase memory times (*T_M_*), a 48 ns π/2–τ– π Gaussian pulse sequence was used with a two-step phase cycling, while τ was increased in 4 ns steps. PELDOR measurements were performed with a Bruker EN5107D2 dielectric resonator at 50 K using a dead-time free four-pulse sequence and a 16-step phase cycling cycling (x[x][x_p_]x) [55,56]. A 38 ns Gaussian pulse (full width at half maximum (FWHM) of 16.1 ns) was used as the pump pulse with a 48 ns (FWHM of 20.4 ns) Gaussian observer pulses. The pump pulse was set to the maximum of the echo-detected field swept spectrum and the observer pulses were set at 80 MHz lower. The deuterium modulations were averaged by increasing the first interpulse delay by 16 ns for 8 steps. The five-pulse PELDOR/DEER experiments were performed according to the pulse sequence π/2_obs_ − (τ/2 – t_0_) − π_pump_ – t_0_ − π_obs_ − t′ − π_pump_ − (τ − t′ + δ) − π_obs_ − (τ_2_ + δ). Experiments were performed at 50 K using 48 ns Gaussian observer pulses and a 16-step phase cycling (xx_p_ [x] [x_p_]x). A 36 ns pump pulse was used at ν_obs_ + 80 MHz. Nuclear modulation averaging was performed analogous to 4-pulse PELDOR (16 ns shift in 8 steps) with a corresponding shift of the standing pump pulse. The four-pulse data analysis was performed using Tikhonov regularization as implemented in the MATLAB-based DeerAnalysis2019 package [57]. The background arising from intermolecular interactions were removed from the primary data V(t)/V(0) and the resulting form factors F(t)/F(0) were fitted with a model-free approach to distance distributions. For an error estimation of the probability distribution, the distances for different background functions were determined through gradually changing the time window and the dimensionality for the spin distribution (see Supplementary Table S2). The data was additionally analyzed to predict the distances (and the background) in a user-independent manner using the deep neural network (DEERNet) analysis, which is hosted by the DeerAnalysis2019 package [58,59]. Samples for which both 4-pulse and 5-pulse data are available were globally analyzed using the Python based DeerLab program [60]. Distance distributions for the structures (PDB 8BJC and 1FD9) were simulated using a rotamer library approach using the MATLAB-based MMM2022.2 software package [58].

### Small angle X-ray scattering (SAXS)

SAXS experiments were carried out at the EMBL-P12 bioSAXS beam line, DESY [61]. SEC-SAXS data were collected [62], *I*(*q*) vs *q*, where *q* = 4*π*sin*q*/*λ* is the scattering angle and λ the X-ray wavelength (0.124 nm; 10 keV). Data collection was carried out at 20 °C using a Superdex200 Increase 5/150 analytical SEC column (GE Healthcare) equilibrated in the appropriate buffers (see Table S3) at flow rates of 0.3 mL/min. Automated sample injection and data collection were controlled using the *BECQUEREL* beam line control software [63]. The SAXS intensities were measured from the continuously-flowing column eluent as a continuous series of 0.25 s individual X-ray exposures, using a Pilatus 6M 2D-area detector for a total of one column volume (ca. 600-3000 frames in total, see Table S3). The radial averaging of the data one-dimensional *I*(*q*) vs *q* profiles was carried out with the SASFLOW pipeline incorporating RADAVER from the ATSAS 2.8 software suite [64]. The individual frames obtained for each SEC-SAXS run were processed using CHROMIXS [65]. Briefly, individual SAXS data frames were selected across the respective sample SEC-elution peaks and appropriate solute-free buffer regions of the elution profile were identified, averaged and then subtracted to obtain individual background-subtracted sample data frames. The radius of gyration (*R_g_*) of each data frame was assessed in CHROMIXS and frames with equivalent *R_g_* were scaled and subsequently averaged to produce the final one-dimensional and background-corrected SAXS profiles. Only those scaled individual SAXS data frames with a consistent *R_g_* through the SEC-elution peak that were also evaluated as statistically similar through the measured *q*-range were included to produce the final SAXS profiles. Corresponding UV traces were not measured; the column eluate was directly moved to the P12 sample exposure unit after the SEC column, forgoing UV absorption measurements, to minimize unwanted band-broadening of the sample. All SAXS data-data comparisons and data-model fits were assessed using the reduced *c*^2^ test and the Correlation Map, or CORMAP, *p*-value [66]. Fits within the *c*^2^ range of 0.9–1.1 or having CORMAP *p*-values higher than the significance threshold cutoff of a = 0.01 are considered excellent, i.e., absence of systematic differences between the data-data or data-model fits at the significance threshold.

Primary SAXS data were analysed using PRIMUS as well as additional modules from the ATSAS 3.0.1 software suite [67]. *R_g_* and the forward scattering at zero angle, *I*(0) were estimated via the Guinier approximation [68] (ln(*I*(*q*)) vs. *q*^2^ for *qR_g_* < 1.3) and the real-space pair distance distribution function, or *p*(*r*) profile (calculated from the indirect inverse Fourier transformation of the data, thus also yielding estimates of the maximum particle dimension, *D_max_*, Porod volume, *V_p_*, shape classification, and concentration-independent molecular weight [69–71]). Dimensionless Kratky plot representations of the SAXS data (*qR* ^2^(*I*(*q*)/*I*(0)) vs. *qR*_g_ ) were generated as previously described [72]. All collected SAXS data are reported in Tables S3.

### Rigid body modeling

Rigid-body normal mode analysis of *Lp*MIP was performed using the program SREFLEX [73] using the *Lp*MIP *apo* and JK095-bound X-ray crystal structures (PDB: 1FD9, 8BJD and 8BJC) as templates. CRYSOL was used to assess data-model fits [74].

### Thermal stability assay

10 µg of purified *Lp*MIP constructs in 20 mM Tris pH 7, 150 mM NaCl were incubated with a final concentration of 0.02 % DMSO or a 5-fold molar excess of JK095 in DMSO (0.02 % final concentration). 2.5 µL of a 50x SYPRO Orange (Merck) stock was added to each sample directly before measurement of the melting temperature in a 96-well plate on a QuantStudio 1 Real-Time PCR System reader (Thermo Fisher) with a temperature increase of 0.05 °C/min. The fluorescence of SYPRO Orange was measured using the filter calibrated for SYBR GREEN with an excitation filter of 470 ± 15 nm and an emission filter of 520 ± 15 nm.

### Data availability

The coordinates of the refined models and structure factors have been deposited into the PDB repository: 8BJC for *Lp*MIP^1-213^, 8BJD for *Lp*MIP^1-213^JK095, 8BK6 for *Lp*MIP^100-213^ JK095, 8BK5 for *Lp*MIP^77-213^ JK095, 8BJE for *Lp*MIP^77-213^ JK236 and 8BK4 for *Tc*MIP JK236. The NMR backbone assignment of *Lp*MIP^100-213^ has been deposited in the BioMagResBank (www.bmrb.io) under the accession number 51861. The NMR backbone assignments for full-length *Lp*MIP^1-213^ and *Lp*MIP^77-213^ are available from the BMRB under the accession numbers 7021 and 6334, respectively [38,75]. SAXS data for full-length *Lp*MIP have been deposited in the SASBDB under the accession numbers SASDSY6 (apo state), SASDSZ6 (with JK095) and SASDS27 (with JK236).

## Author contributions

Sample preparation: CW, VHPC, FT, BG; Biochemistry: VHCP, FT, BG; X-ray crystallography: JJW, BG, AG; NMR spectroscopy: CW, VHPC, FT, BG; EPR spectroscopy: VHPC, MD, BJ; SAXS: CW, FT, BG, JMH; Inhibitor synthesis: PK; Conceptualization: UAH; Funding acquisition: BJ, FH, AG, UAH; Supervision: BG, BJ, FH, AG, UAH; Paper writing – first draft: UAH; Paper writing – review and editing: CW, JJW, BG, BJ, AG, UAH; visualization: CW, JJW, VHPC, MD, BG, UAH. All authors read and approved the final version of the manuscript.

## Acknowledgments

We thank Sarah-Ana Mitrovic, Hannah Niederlechner, Sabine Häfner and Dania Rose-Sperling for technical assistance and Robin Deutscher for support with synthesis. VHPC acknowledges a <u>DAAD-</u> <u>CONACYT PhD fellowship. BG</u> acknowledges a PhD fellowship by the Max Planck Graduate Center (MPGC). Access to beamline P12, DESY, Hamburg was made available via iNEXT-ERIC (BAG proposal #SAXS-1106 (to UAH)). We are grateful to Shibom Basu and Montserrat Soler Lopez at the ESRF for providing assistance at beamlines ID23-2 and ID30A-3 (BAG proposals #MX-2268 and #MX- 2407 to AG). We thank Andreas Schlundt for organizational support with SAXS measurements. We thank the Centre of Biomolecular Magnetic Resonance (BMRZ) at the Goethe University Frankfurt funded by the state of Hesse and the Jena School for Microbial Communication (JSMC) for support. Funded by the Federal Ministry of Education and Research (BMBF) project iMIP (16GW0211 to FH, 16GW0214 to UAH). BJ acknowledges financial support through the Emmy Noether program (JO 1428/1−1) and a large equipment funding (438280639) from the Deutsche Forschungsgemeinschaft (DFG). Supported by the DFG under Germany’s Excellence Strategy - EXC 2051 - Project ID 390713860 and the collaborative research cluster SFB1127/3 ChemBioSys— Project ID 239748522 (to UAH). UAH acknowledges an instrumentation grant by the REACT-EU EFRE Thuringia (Recovery assistance for cohesion and the territories of Europe, European Fonds for Regional Development, Thuringia) initiative of the European Union.

---

**Scheme 1:**
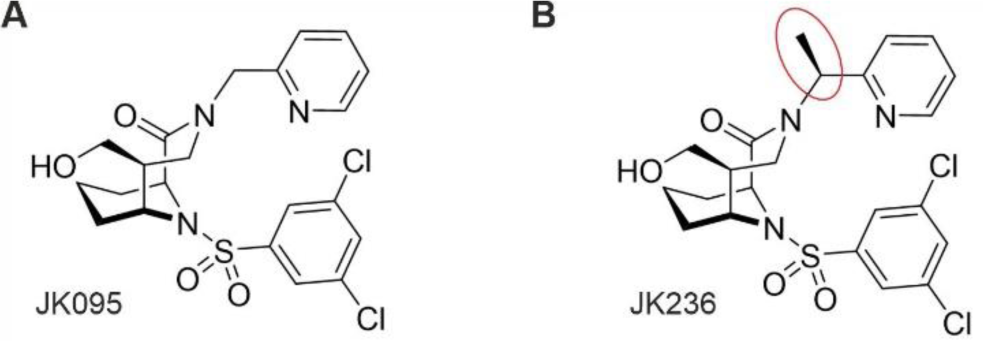
[**4.3.1]-aza-bicyclic sulfonamide inhibitors used in this study.** JK095 (**A**) and JK236 (**B**) differ by the insertion of a stereospecific methyl group in the pyridine linker.

