## Supporting information for "*Legionella pneumophila* macrophage infectivity potentiator protein appendage domains modulate protein dynamics and inhibitor binding"

**Table S1: Data collection and refinement statistics (molecular replacement)**

| | *Lp*MIP$^{1-213}$ | *Lp*MIP$^{1-213}$ + JK095 | *Lp*MIP$^{100-213}$ + JK095 | *Lp*MIP$^{77-213}$ + JK095 | *Lp*MIP$^{77-213}$ + JK236 | *Tc*MIP + JK236 |
|---|---|---|---|---|---|---|
| **PDB-ID** | 8BJC | 8BJD | 8BK6 | 8BK5 | 8BJE | 8BK4 |
| Wavelength | 0.9763 | 0.9763 | 0.9763 | 0.9763 | 0.9763 | 0.9795 |
| Resolution range | 62.24 - 1.71 (1.772 - 1.71) | 48.07 - 2.4 (2.486 - 2.4) | 59.92 - 2.263 (2.344 - 2.263) | 29.7 - 1.44 (1.492 - 1.44) | 45.86 - 1.491 (1.544 - 1.49) | 35.99 - 1.342 (1.39 - 1.342) |
| Space group | P 43 21 2 | P 43 21 2 | P 43 21 2 | P 31 2 1 | P 31 2 1 | P 21 21 21 |
| *a*, *b*, *c* (Å) | 77.773 77.773 103.789 | 76.752 76.752 103.597 | 73.557 73.557 103.286 | 53.54 53.54 77.36 | 52.951 52.951 73.146 | 42.493 57.529 67.712 |
| α, β, γ (°) | 90, 90, 90 | 90, 90, 90 | 90, 90, 90 | 90, 90, 120 | 90, 90, 120 | 90, 90, 90 |
| Unique reflections | 19488 (241) | 12644 (1218) | 13801 (1344) | 23619 (2330) | 15279 (250) | 28364 (209) |
| Completeness (%) | 55.53 (7.00) | 99.91 (99.75) | 99.63 (99.18) | 99.21 (98.69) | 76.74 (12.73) | 74.79 (5.62) |
| Mean I/sigma(I) | 12.2(2.61) | 11.1(2.10) | 8.0(2.32) | 9.92(2.59) | 15.33(2.04) | 15.56(1.30) |
| Wilson B-factor | 22.40 | 45.77 | 48.51 | 18.67 | 19.02 | 13.38 |
| R-meas | 0.09 | 0.06 | 0.06 | 0.07 | 0.08 | 0.08 |
| CC1/2 | 93.3(24.0) | 99.7(22.1) | 98.5(32.3) | 99.6(21.8) | 99.1(27.7) | 99.2(26.0) |
| Reflections used in refinement | 19488 (241) | 12642 (1218) | 13760 (1334) | 23616 (2330) | 15273 (250) | 28314 (209) |
| Reflections used for R-free | 971 (16) | 607 (56) | 663 (53) | 1137 (132) | 745 (15) | 1444 (10) |
| R-work | 0.2455 (0.3083) | 0.2357 (0.3254) | 0.2868 (0.3346) | 0.1838 (0.3227) | 0.1975 (0.2236) | 0.1968 (0.2986) |
| R-free | 0.2943 (0.3324) | 0.2898 (0.4187) | 0.3482 (0.4109) | 0.2227 (0.3722) | 0.2332 (0.3624) | 0.2205 (0.2814) |
| Number of non-hydrogen atoms | 1742 | 1644 | 1737 | 1271 | 1122 | 1507 |
| macromolecules | 1554 | 1573 | 1684 | 1059 | 1020 | 1316 |
| ligands | 105 | 72 | 40 | 110 | 85 | 43 |
| solvent | 139 | 20 | 13 | 148 | 58 | 148 |
| Protein residues | 205 | 208 | 226 | 135 | 134 | 163 |
| RMS(bonds) | 0.006 | 0.008 | 0.01 | 0.009 | 0.017 | 0.006 |
| RMS(angles) | 0.96 | 1.14 | 1.54 | 1.11 | 1.69 | 0.92 |
| Ramachandran favored (%) | 98.03 | 95.63 | 97.30 | 97.74 | 96.21 | 99.38 |
| Ramachandran allowed (%) | 1.97 | 4.37 | 2.25 | 2.26 | 3.03 | 0.62 |
| Ramachandran outliers (%) | 0 | 0 | 0.45 | 0 | 0.76 | 0 |
| Rotamer outliers (%) | 0.59 | 1.16 | 4.30 | 0.84 | 0.88 | 0.72 |
| Clashscore | 6.78 | 7.35 | 9.15 | 3.53 | 5.62 | 4.10 |
| Average B-factor | 27.24 | 46.16 | 53.96 | 24.55 | 31.56 | 17.53 |
| macromolecules | 26.94 | 45.64 | 53.95 | 23.63 | 31.19 | 16.61 |
| ligands | 28.82 | 61.86 | 55.20 | 16.76 | 29.50 | 16.37 |
| solvent | 29.95 | 47.15 | 50.25 | 34.46 | 39.63 | 26.06 |

*Values in parentheses are for the highest resolution shell.

**Table S2:** Parameters for error estimation of the probability distributions obtained using Tikhonov regularization for full-length *Lp*MIP K80C. Validation was performed as featured in the DeerAnalysis2019 software package[1].

| **Sample** | **Error validation** | | | | | **Regularization parameter (α)** |
|---|---|---|---|---|---|---|
| | dimensionality (*d*) | | starting time window | | | |
| | range | steps | range (ns) | steps | $T_{max}$ (µs) | |
| **K80C** | 2.8-3.2 | 9 | 240-1000 | 11 | 5.36 | 1258 |
| **K80C + JK095** | 2.8-3.2 | 9 | 240-1000 | 11 | 4.86 | 1584 |
| **K80C + JK236** | 2.8-3.2 | 9 | 240-1000 | 11 | 4.38 | 1000 |

**Table S3:** SAXS data reporting table for full-length *Lp*MIP.

| **Sample details** | | | |
|---|---|---|---|
| SAMPLE | *Lp*MIP *apo* | *Lp*MIP + JK095 | *Lp*MIP + JK236 |
| SASBDB Accession Codes | SASDSY6 | SASDSZ6 | SASDS27 |
| Organism | | *Legionella pneumophila* | |
| NCBI protein accession ID | | 66489975 | |
| (amino acid range) | | 1-213* | |
| SEC-SAXS buffer | | 20 mM Tris pH 7.5, 10 mM DTT | |
| NaCl concentration | | 150 mM | |
| Sample injection volume | | 45 | |
| Sample injection conc. | | 10 mg/mL | |
| SEC column | | S200 Increase 5/150 | |
| SEC flow rate | | 0.3 ml/min | |
| SEC temperature | | 20 ºC | |
| **Instrument details** | | | |
| Instrument | | EMBL P12 bioSAXS beam line, DESY, Hamburg | |
| Exposure time/# frames | | 0.25 s (2400) | |
| X-ray wavelength/energy | | 0.124 nm (9996.5 eV) | |
| Sample-to-detector distance | | 3 m | |
| Scattering intensity scale | | Arbitrary unit, a.u. | |
| SEC-SAXS primary data processing | | CHROMIXS (ATSAS 3.0.1) | |
| # frames used for averaging | 90 | 78 | 77 |
| Working s-range ($nm^{-1}$) | 0.03-7.43 | 0.03-7.43 | 0.03-7.43 |
| **Guinier analysis:** | | | |
| Primary data analysis software | | PRIMUS (ATSAS 3.0.1) | |
| Guinier I(0) (σ) | 8491(8) | 8031(10) | 7835(8) |
| $R_g$ (Guinier, nm) (σ) | 3.13(0.01) | 3.11(0.01) | 3.04(0.01) |
| $sR_g$ range | 0.34-1.30 | 0.41-1.29 | 0.27-1.30 |
| **p(r) analysis:** | | | |
| Method | | GNOM 5 | |
| I(0), POR (σ) | 8526(7) | 8068(9) | 7686(8) |
| $R_g$ (POR, nm) (σ) | 3.17(0.1) | 3.14(0.01) | 3.07(0.01) |
| $D_{max}$ (nm) | 10.0 | 10.0 | 9.4 |
| Quality of fit, CorMap P / $\chi^2$ | 2.75E-04 / 1.04 | 6.7E-0.2 / 1.02 | 1.7E-0.2 / 1.00 |
| Porod volume ($nm^3$) | 66 | 62 | 63 |
| Shape classification | flat | flat | flat |
| **Molecular Weight analysis:** | | | |
| MW, calculated from amino acid sequence, kDa | 45.6 (dimer) | 46.5 (dimer with 2x JK095) | 46.5 (dimer with 2x JK236) |
| MW from SAXS data, kDa | 41-46 | 44-48 | 46-49 |
| **Rigid body/Normal mode modelling:** | | | |
| Method | | SREFLEX (five individual fits) | |
| Symmetry | P1 | P1 | P1 |
| Template | 8BJC | 8BJD | 8BJC |
| Initial template fit, CorMap P / $\chi^2$ | 3.10E-59 / 14.30 | 1.87E-63 / 16.28 | 7.73E-60 / 10.45 |
| Final model fit, CorMap P / $\chi^2$ | 6.76E-03 / 1.16 | 5.3E-05 / 1.30 | 2E-06 / 1.75 |
| | 2.56E-08 / 1.26 | 1.3E-05 / 1.37 | 7.99E-10 / 1.86 |
| | 1.24E-11 / 1.32 | 4.11E-07 / 1.38 | 7E-06 / 1.88 |
| | 4.23E-04 / 1.32 | 7.99E-10 / 1.39 | 8.22E-07 / 1.90 |
| | 1.60E-09 / 1.33 | 7.99E-10 / 1.42 | 4.11E-07 / 1.93 |

*The nomenclature for *Lp*MIP$^{1-213}$ used in this manuscript refers to the processed protein after cleavage of the N-terminal signal peptide comprising residues 1-24 according to GenBank entry AAB22717.1. (https://www.ncbi.nlm.nih.gov/protein/AAB22717.1)

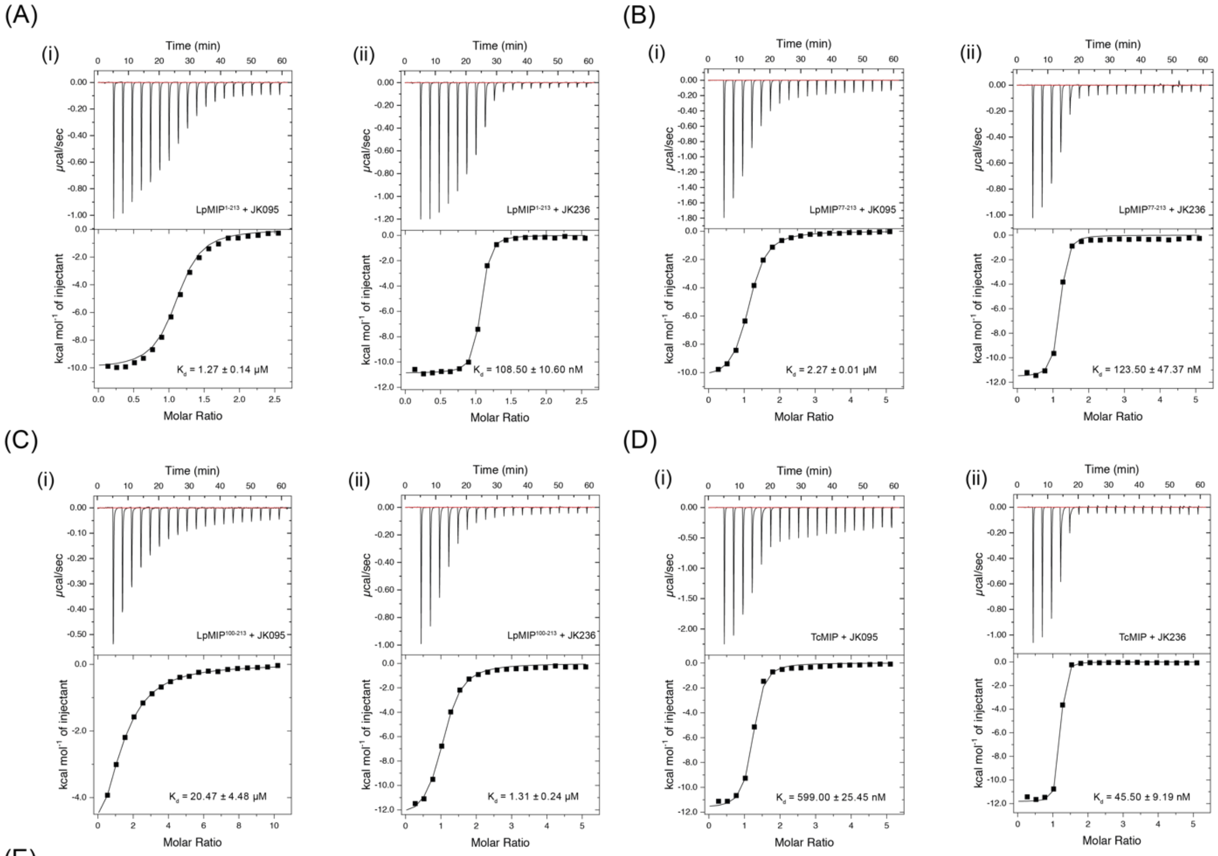


(E)

| | $LpMIP^{1-213}$ | | $LpMIP^{77-213}$ | | $LpMIP^{100-213}$ | | TcMIP | |
|---|---|---|---|---|---|---|---|---|
| | + JK095 | + JK236 | + JK095 | + JK236 | + JK095 | + JK236 | + JK095 | + JK236 |
| Kd | 1.27 ± 0.14 µM | 108.50 ± 10.60 nM | 2.27 ± 0.01 µM | 123.50 ± 47.37 nM | 20.47 ± 4.48 µM | 1.31 ± 0.24 µM | 599.00 ± 25.45 nM | 45.50 ± 9.19 nM |
| ΔH (cal/mol) | -10770 ± 438 | -11185 ± 431 | -9886 ± 953 | -11755 ± 275 | -8968 ± 1769 | -12590 ± 480 | -11225 ± 289 | -11795 ± 289 |
| ΔS (cal/mol/deg) | -9.15 ± 1.20 | -5.65 ± 1.25 | -7.33 ± 3.20 | -7.75 ± 0.14 | -8.60 ± 6.36 | -15.3 ± 1.97 | -9.18 ± 1.06 | -5.96 ± 1.38 |
| ΔG (cal/mol) | -8043 ± 80 | -9501 ± 56 | -7700 ± 13 | -9445 ± 233 | -6405 ± 126 | -8030 ± 109 | -8489 ± 26 | -10018 ± 123 |
| N | 0.97 ± 0.17 | 0.99 ± 0.03 | 1.17 ± 0.14 | 1.02 ± 0.08 | 1.19 ± 0.02 | 0.97 ± 0.04 | 1.13 ± 0.01 | 0.99 ± 0.12 |

**Fig. S1: Isothermal titration calorimetry of *Legionella pneumophila* and *Trypanosoma cruzi* MIP with bicyclic inhibitors. (A)** *Lp*MIP$^{1-213}$ (full-length protein), **(B)** *Lp*MIP$^{77-213}$ and (**C**) *Lp*MIP$^{100-213}$ with JK095 and JK236 (i and ii, respectively). (**D**) *Tc*MIP with JK095 (i) and JK236 (ii). Representative ITC traces are shown, all measurements n=2. (**E**) Fitting parameters for ITC measurements.

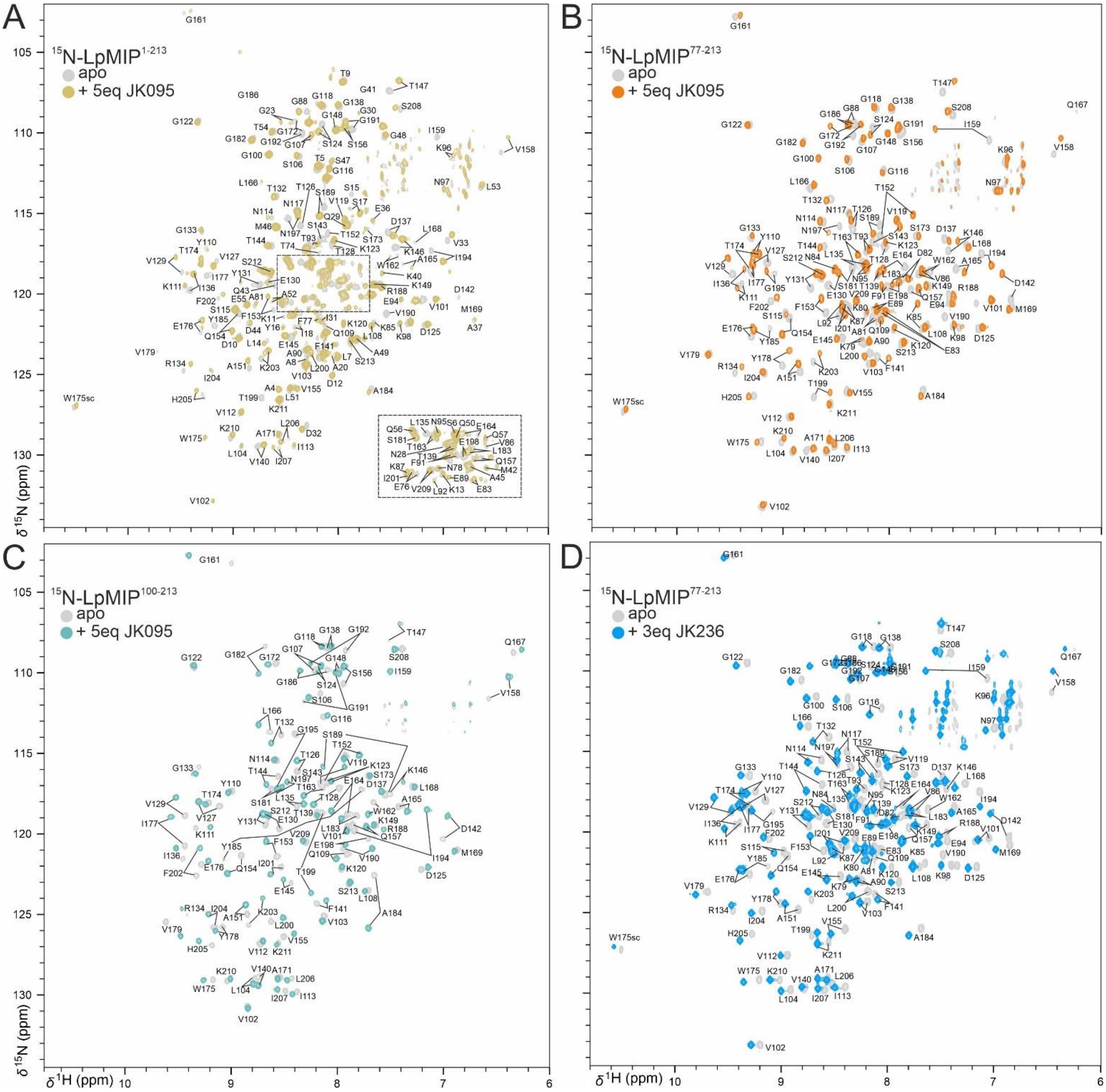


**Fig. S2: Backbone NMR assignments of *Lp*MIP constructs in the apo and inhibitor bound states.** Backbone assignments of (**A**) full-length *Lp*MIP without (grey) and with JK095 (sand), (**B**) *Lp*MIP$^{77\text{-}213}$ without (grey) and with JK095 (orange), (**C**) *Lp*MIP$^{100\text{-}213}$ without (grey) and with JK095 (teal), and (**D**) *Lp*MIP$^{77\text{-}213}$ without (grey) and with JK236 (blue). Previously published backbone amide resonance assignments for full-length *Lp*MIP (BMRB entry 7021) and *Lp*MIP$^{77\text{-}213}$ (BMRB entry 6334) could be partially transferred to our spectra and were verified using 3D assignment experiments under our buffer conditions. In contrast, the assignment of *Lp*MIP$^{100\text{-}213}$ had to be determined *de novo*, as the $^{1}$H, $^{15}$N-HSCQ spectrum of this construct differed significantly from the resonances of the FKBP domain in both *Lp*MIP$^{77\text{-}213}$ and full-length *Lp*MIP. The assignment for *Lp*MIP$^{100\text{-}213}$ has been deposited in the BMRB under accession number 51861.

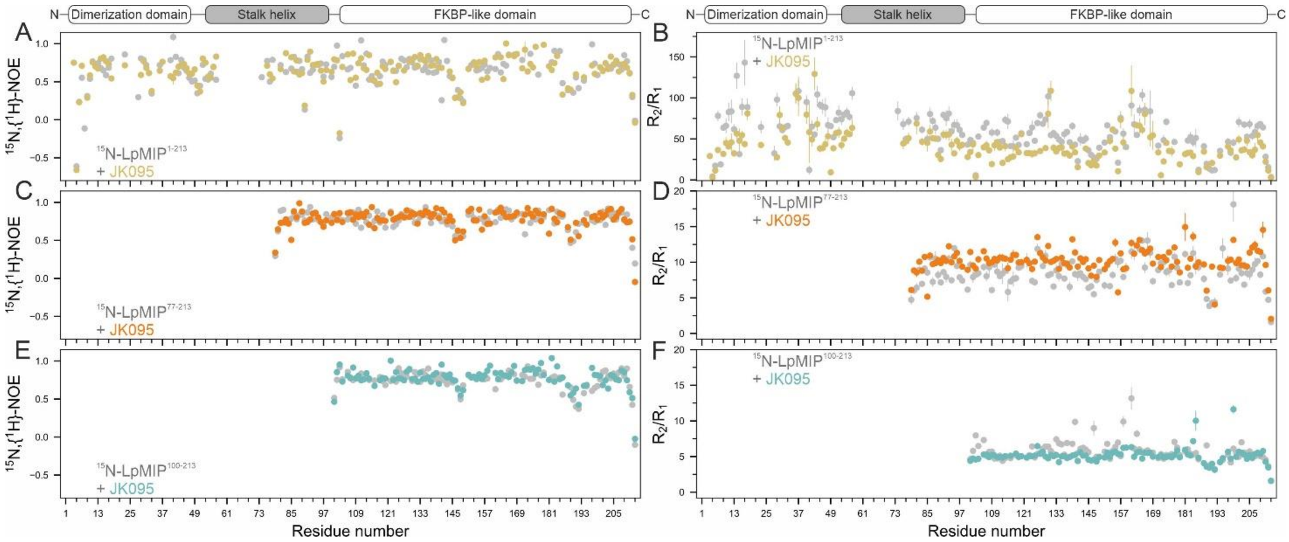


**Fig. S3: Fast backbone dynamics of *Lp*MIP constructs in the absence and presence of JK095.** $^{15}$N, *{$^{1}$H}-NOE* and $R_1/R_2$ relaxation measurements of full-length *Lp*MIP (A, B), *Lp*MIP$^{77-213}$ (C, D) and *Lp*MIP$^{100-213}$ (E, F) without (grey circles) or in the presence of a five-fold molar excess of JK095 (coloured circles). More flexible regions are the loops connecting β3a/β3b as well β4/β5 loop centered around residues ~145 and ~190, respectively.

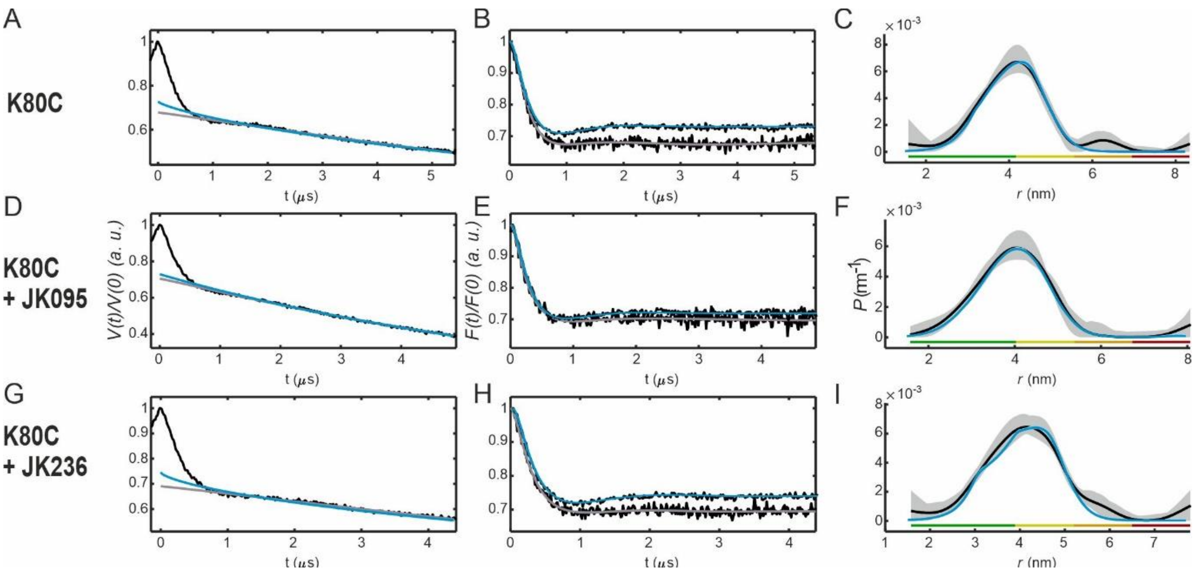


**Fig. S4: PELDOR/DEER data analysis for *Lp*MIP K80C with the inhibitors JK095 and JK236.** (A, D, G) The primary data (black) overlaid with the intermolecular (background) contribution from deep neural network analysis (blue) and Tikhonov regularization (grey). (B, E, H) The background corrected form factors overlaid with the fits. (C, F, I) The corresponding distance distributions.

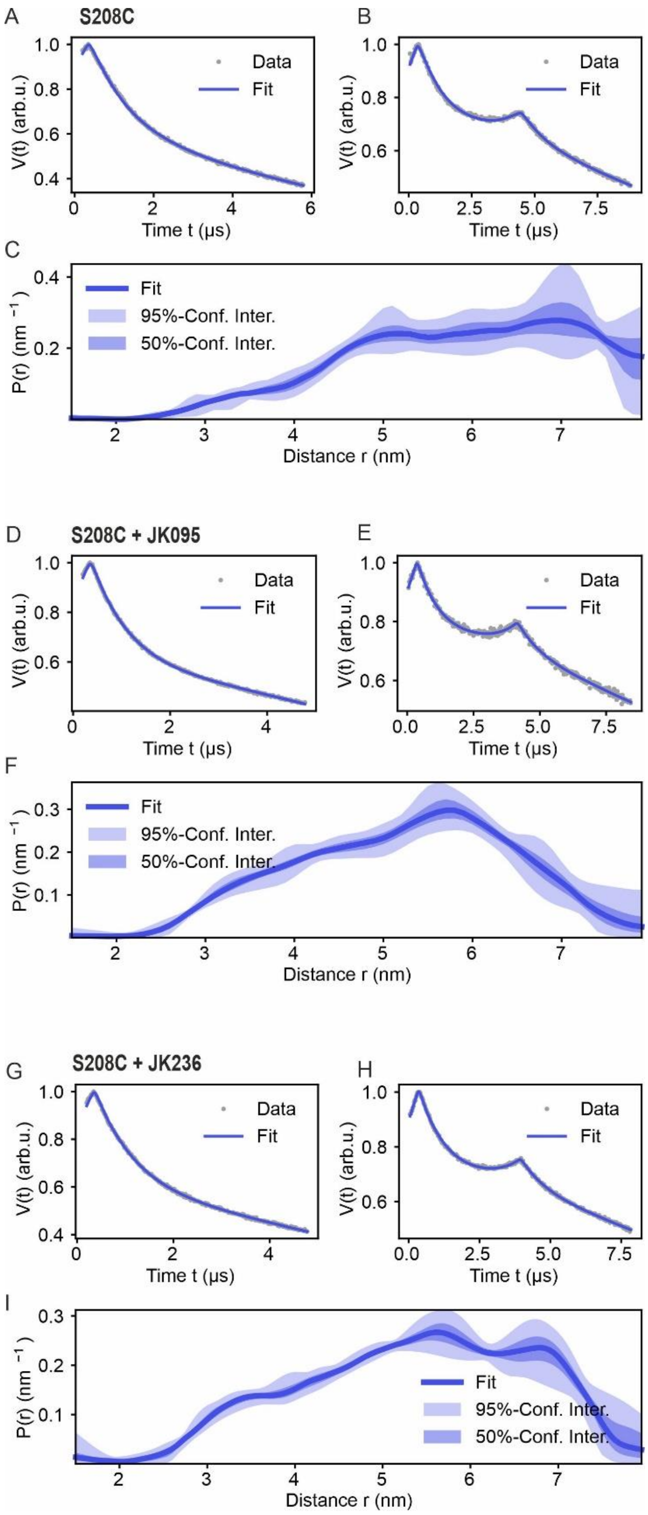


**Fig. S5: PELDOR/DEER data analysis for *Lp*MIP S208C with the inhibitors JK095 and JK236.** The 4-pulse and 5-pulse PELDOR data were globally analysed using the Python based DeerLab program. (A, D, G) The 4-pulse PELDOR data (grey) overlaid with the fit (blue). (B, E, H) The 5-pulse PELDOR data (grey) overlaid with the fit (blue); (C, F, I) The corresponding distance distributions with 50%- (shaded in dark blue) and 95% confidence intervals (shaded in light blue).

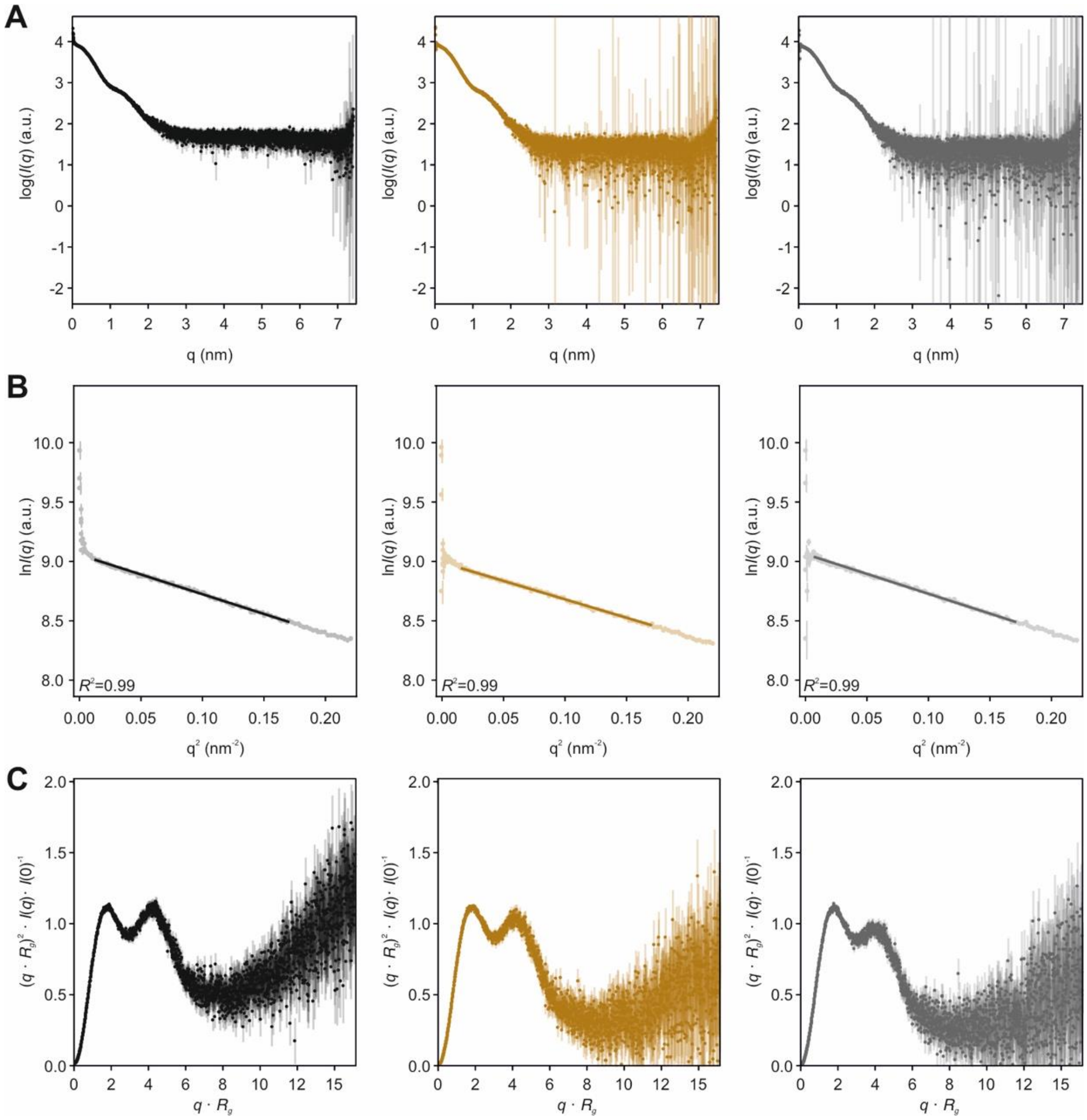


**Fig. S6 – SAXS data collection of *Lp*MIP.** (A) X-ray scattering profiles of *Lp*MIP apo (black), *Lp*MIP + JK095 (brown), and *Lp*MIP + JK236 (grey) plotted as the logarithm of the scattering intensity log($I(q)$) (a.u., arbitrary units) versus the momentum transfer, $q$. (B) Guinier-plots (ln$I(q)$ vs $q^2$, plotted to low-angle: $qR_g <$ 1.3) of *Lp*MIP apo (black), *Lp*MIP + JK095 (brown), and *Lp*MIP + JK236 (grey). (C) Dimensionless Kratky-plots for *Lp*MIP apo (black), *Lp*MIP + JK095 (brown), and *Lp*MIP + JK236 (grey) plotted as $(qR_g)^2I(q)I(0)^{-1}$ vs. $qR_g$..

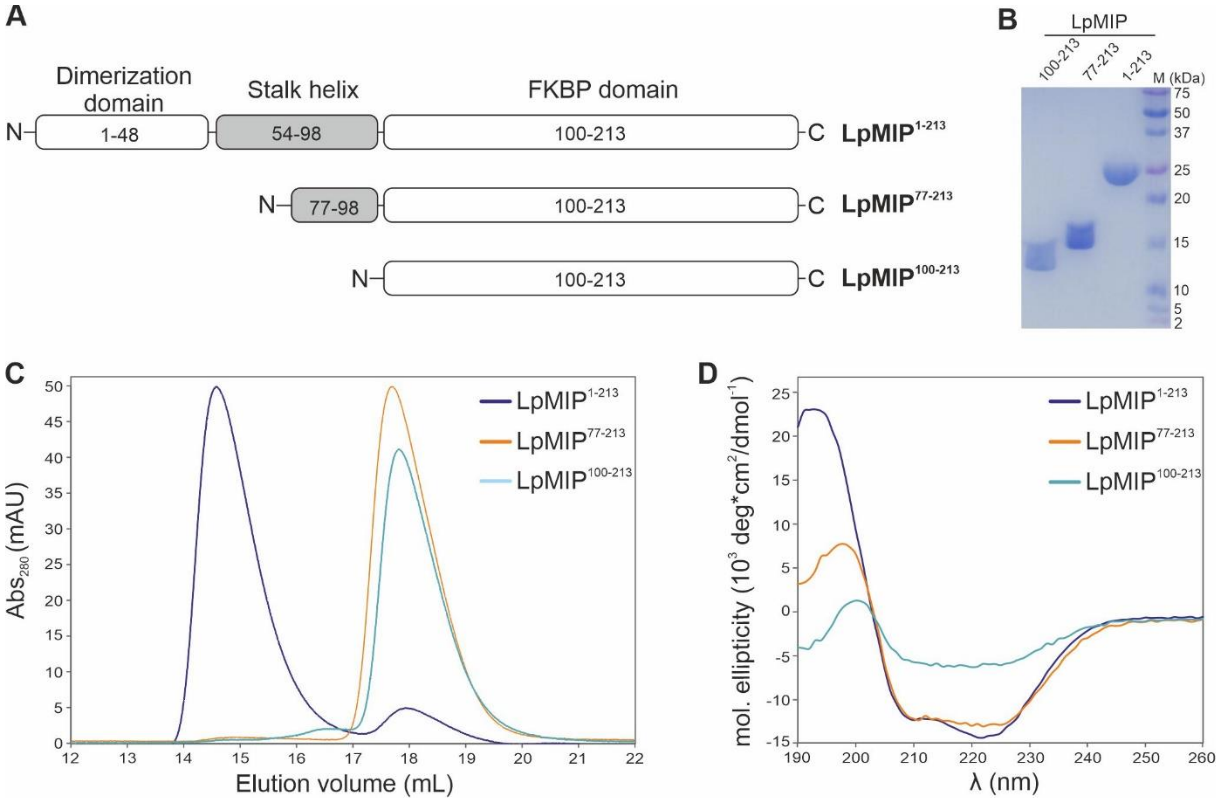


**Fig. S7: Purification and structural integrity of *Legionella pneumophila* MIP deletion constructs**. (**A**) Schematic of *Lp*MIP constructs used in this study. (**B**) SDS-PAGE of full-length *Lp*MIP (residues 1-213) and N-terminally truncated versions lacking the dimerization domain and half the stalk helix (*Lp*MIP$^{77-213}$) or the dimerization domain and the entire stalk helix (*Lp*MIP$^{100-213}$). (**C**) Analytical size exclusion chromatography of the three *Lp*MIP constructs. Note that *Lp*MIP$^{1-213}$ forms a dimer, while the two shorter constructs are monomeric. (**D**) Circular dichroism spectra of the three *Lp*MIP constructs displays the expected secondary structure content.

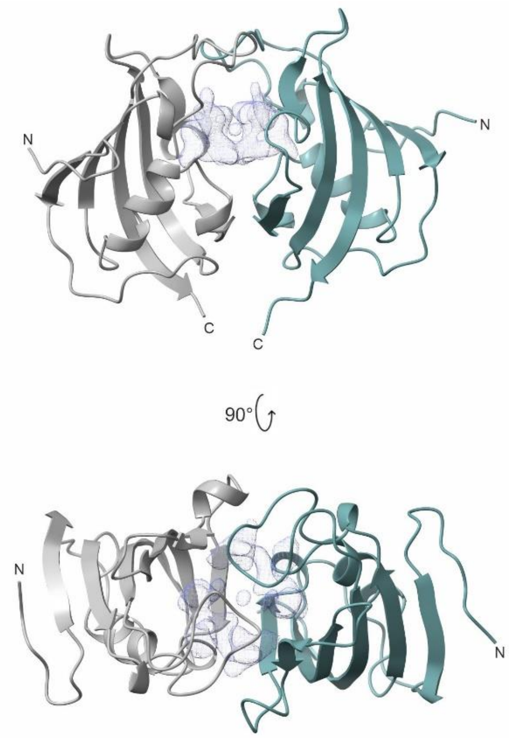


**Fig. S8: *Lp*MIP$^{100-213}$ crystallizes as a dimer.** In the crystal structure of *Lp*MIP$^{100-213}$ (PDB: 8BK6), the two protomers (grey, dark teal) in the unit cell align with an RMSD of 0.327 Å. The JK095 binding site cannot be defined clearly, instead there is density throughout the interface (light grey mesh). Of note, NMR spectroscopically derived rotation correlation times indicate that in solution, *Lp*MIP$^{100-213}$ is monomeric, irrespective whether JK095 is present or not (see main text for details).

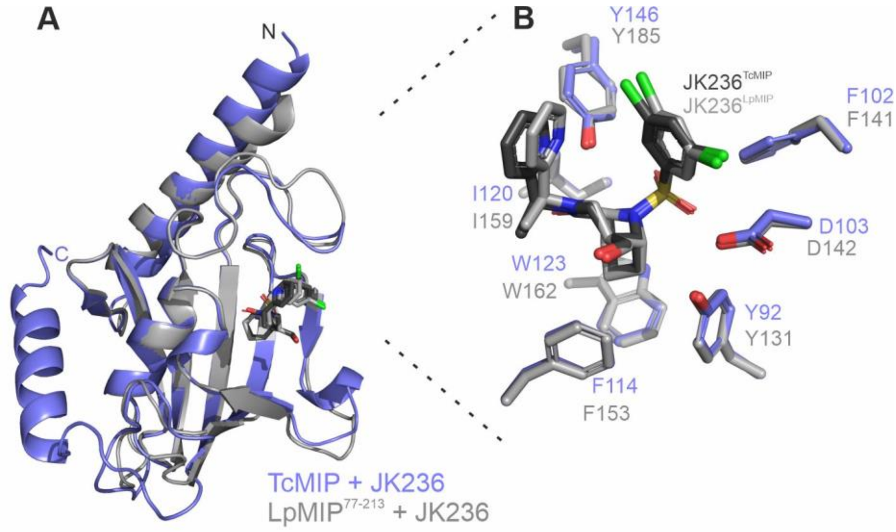


**Fig. S9: Comparison of *Trypanosoma cruzi* and *Legionalla pneumophila* MIP in complex with a [4.3.1]-aza-bicyclic sulfonamide inhibitor.** (**A**) Overlay of the X-ray structures of *Tc*MIP (blue) and *Lp*MIP$^{77\text{-}213}$ (grey) in complex with JK236 (PDB-IDs: 8BJE, 8BK4). Both structures align with a backbone RMSD of 0.51 Å. (**B**) Zoom into the binding site of *Tc*MIP and *Lp*MIP bound to JK236. Between JK236-bound *Tc*MIP and *Lp*MIP$^{77\text{-}213}$, the inhibitor binding stance is nearly identical and only a single rotamer of the hydroxymethyl group was observed. Likewise, the inhibitor's pyridine-linker methyl-group was solvent exposed in both proteins.

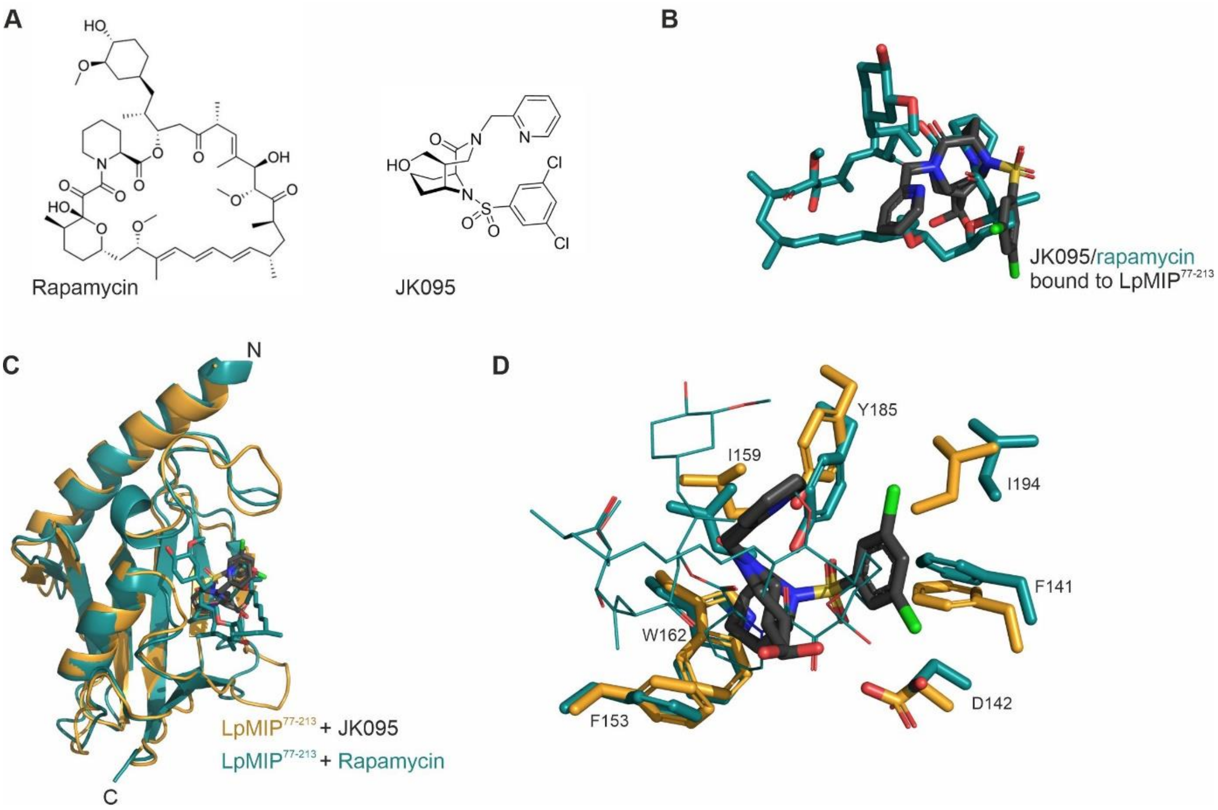


**Fig. S10: Comparison of *Lp*MIP$^{77\text{-}213}$ bound to rapamycin or a bicyclic sulfonamide inhibitor.** (**A**) Structures of rapamycin and JK095. (**B**) Overlay of rapamycin (dark teal) and JK095 (grey) bound to *Lp*MIP$^{77\text{-}213}$ (PDB IDs: 2VCD, 8BK5). (**C**) Overlay of *Lp*MIP$^{77\text{-}213}$ in complex with rapamycin (dark teal) or JK095 (orange, ligand shown in dark grey). (**D**) Zoom into the ligand binding site, residues important for ligand contacts are shown as sticks. For better visualization, rapamycin is shown with thin lines.
